# Orexinergic projections to substantia innominata mediate arousal and analgesia

**DOI:** 10.1101/2024.10.29.620973

**Authors:** Xuaner Xiang, Fei Wang, Chao Chen, Zhonghui Guan, Wei Zhou

## Abstract

Understanding neural circuits involved in anesthesia is crucial for improving its safety and efficacy. Hypothalamic orexin neurons (LHA^OX^), projecting broadly, are essential in regulating arousal and pain. However, the precise targets remain unclear. Here we investigated the orexin projections to the substantia innominata (SI). Combining optogenetics, fiber photometry, and EEG/EMG allowed us to manipulate orexin activities, while simultaneously recording local ligand release and global cortical activities during anesthesia. Brain slice electrophysiology revealed the synaptic connections in the SI, while RNAscope was employed to examine the distribution of orexin receptors and downstream neuronal types. Presynaptic vesicles were identified in the orexin terminals in the SI, where 49.16% of cells expressed OX2R and 6.8% expressed OX1R. Orexin release in the SI was reversibly suppressed by isoflurane. Optogenetic activation of the LHA^OX^→SI circuit significantly increased orexin release and promoted arousal from various anesthesia stages, including reanimation during 0.75% isoflurane (p < 0.0001), prolongation of 3% isoflurane induction (p = 0.0033), and acceleration of emergence from 2% isoflurane (p < 0.0001). Furthermore, activating this circuit induced analgesia to both thermal (p = 0.0074) and inflammatory (p = 0.0127) pain. Patch-clamp recordings revealed that optogenetic activation of orexin terminals in the SI elicited excitatory postsynaptic currents, which were blocked by the OX2R antagonist. SI contains more GABAergic (28.17%) and glutamatergic (11.96%) neurons than cholinergic neurons (4.13%), all of which expressed OX2R. Thus, LHA^OX^ neurons innervate SI neurons to regulate both arousal and pain predominantly through OX2R.

## Introduction

Anesthesia has been practiced for close to 200 years, yet the underlying neural circuitry remains incompletely understood. Lack of deep understanding leads to adverse events due to unintentional anesthetics overdosing, including neurotoxicity in both developing and aging brains (1), sympathetic nervous system suppression (2), and prolonged postoperative recovery (3). Neural circuits underpin the core elements of anesthesia - hypnosis, amnesia, immobility, and analgesia (4). Among these circuits, the hypothalamic orexin/hypocretin neurons, located in the lateral hypothalamic area (LHA), broadly project throughout the brain and spinal cord, playing essential roles in regulating arousal, feeding, pain perception, stress, reward, and memory formation (5–10).

Orexin neurons release two homologous neuropeptides orexin A and B, which target two G-protein coupled receptors OX1R and OX2R (5,6). Dysfunction of the orexin circuit is linked to narcolepsy with cataplexy, characterized by disrupted sleep-wake patterns and sudden loss of muscle tone (11,12). Studies have shown that OX2R knockout mice exhibit symptoms of narcolepsy, while OX1R knockout mice have been grossly normal, highlighting the critical role of OX2R in sleep-wake control (13). However, locus coeruleus (LC), rich in OX1R, is well-known for its adrenergic neurons and their wake-promoting role. Orexin projections serve diverse functions, for example, the projection to the lateral habenula is involved in regulating aggressive behavior in male mice (14), while the projection to the ventral tegmental area (VTA) has a role in regulating stress-induced cocaine recurrence (15). Animal experiments showed that orexin is involved in various aspects of anesthesia, including emergence, airway patency, autonomic tone, and gastroenteric motility (16–19). The precise orexin circuits responsible for arousal from anesthesia are still not fully understood.

Beyond arousal control, orexin also plays a crucial role in modulating analgesia, an essential component of anesthesia (20,21). Given its widespread projections throughout the brain and spinal cord, orexin contributes to pain modulation at multiple levels. At the spinal cord, orexin directly acts on the dorsal root ganglia to regulate pain signal transmission (22). In the brain, orexin regulates pain via key regions such as the rostral ventromedial medulla (RVM) (23), VTA (24), and periaqueductal gray (PAG) (25), involving various neurotransmitters including dopamine, adenosine, endocannabinoid, and opioid. Nevertheless, the specific circuitry mechanism involving orexin neurons to relieve pain is yet to be determined.

Previously, we mapped the 3D projections of orexin neurons in the whole mouse brain, revealing dense projections to the substantia innominata (SI) (26), located at the anteromedial and ventral part of the cerebral hemisphere. SI is involved in aggressive response, cortical processing, and sleep (27–29). It contains the nucleus basalis of Meynert which is the major source of cholinergic input to the cerebral cortex. Cholinergic output has been known to be a key for cortical activation and anesthesia arousal (30,31). Additionally, SI contains a significant population of GABAergic and glutamatergic neurons (32). Beyond arousal control, SI is part of an affective pain circuit and receives innervation from the central amygdala (CeL) to modulate fear-related behaviors (33). Given that the LHA responds to noxious stimuli and regulates pain processing (34,35), important questions persist regarding the synaptic connections between LHA^OX^ and SI, the distribution of two orexin receptors, and the profile of SI neuronal types innervated by orexin. In this study, we employed optogenetics, EEG/EMG, in vivo fiber photometry, brain slice electrophysiology, and RNAscope in situ hybridization to investigate the LHA^OX^→SI circuit, and its role in arousal and pain regulation.

## Results

### Hypothalamic Orexin Neurons Project to the SI

Our previous results revealed dense fibers observed in the SI (26). To confirm the presence of orexin terminals, we utilized the synaptic vesicle marker - synaptophysin (Syp) - a synaptic vesicle protein. We injected adeno-associated virus (AAV)-FLEX-tdTomato-T2A-SypEGFP into the LHA of the Orexin-Cre mice (Fig. 1A). The T2A sequence encodes a self-cleaving peptide that separates SypEGFP from tdTomato. This process targets SypEGFP to presynaptic vesicles at the nerve terminals, while tdTomato is expressed throughout the cells. Our results showed punctate SypEGFP expression in both the LHA and SI (Fig. 1B), suggesting the presence of the orexin synaptic terminals in these regions.

**Figure 1.**
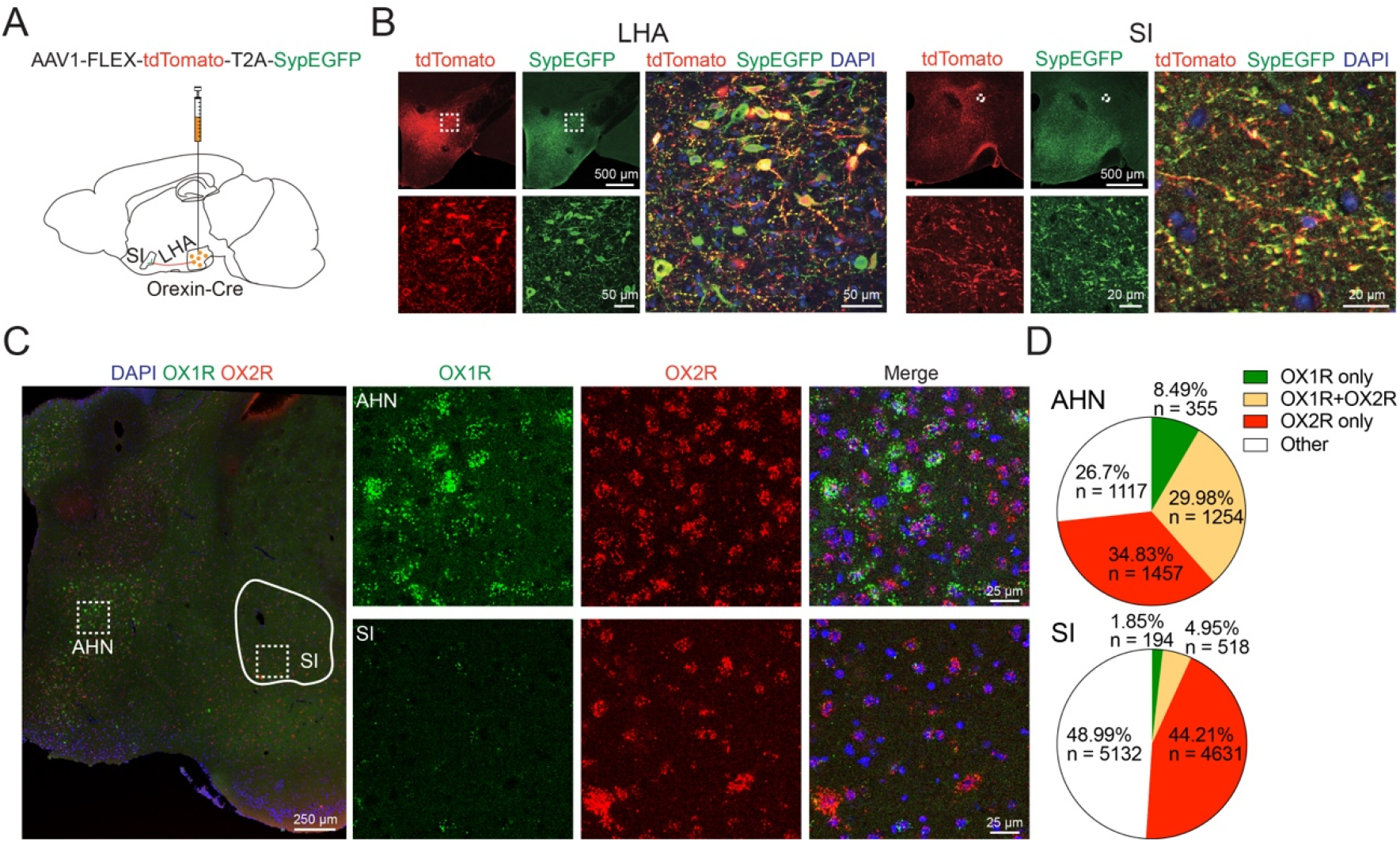
Hypothalamic orexin neurons project to the SI. **A**. *AAV1-FLEX-tdTomato-T2A-SypEGFP* was injected into the lateral hypothalamic area (LHA) of *Orexin-Cre* mice. **B**. Orexin cell bodies in the LHA and fiber terminals in the substantia innominata (SI) are visualized by tdTomato and synaptophysin-EGFP (SypEGFP). **C**. Images of RNAscope in situ hybridization in the AHN and SI. **D**. Distributions of OX1R and OX2R in AHN and SI. AHN, 6 slices from 4 brains. SI, 10 slices from 5 brains. LHA, lateral hypothalamic area; SI, Substantia innominata; AHN, Anterior hypothalamic nucleus.

Then we focused on the distribution of postsynaptic orexin receptors in the SI to further confirm the innervation by the orexin terminals. We used RNAscope in situ hybridization to probe the brain slices for mRNAs of OX1R and OX2R. The results revealed 44.21% of SI cells expressing OX2R only, 1.85% OX1R only, and 4.95% cells expressing both receptors (Fig. 1C, D). The distribution of OX1R and OX2R are highly variable in the brain. As positive controls, it is known that the paraventricular nucleus of the thalamus (PVT) expresses both OX1R and OX2R, while the locus coeruleus (LC) primarily expresses OX1R (9,10). Our RNAscope data confirmed this by showing the even staining of OX1R and OX2R mRNA in the PVT and predominantly OX1R mRNA in the LC. (Supplementary Fig. 1). Additionally, on the same brain slice in the nearby anterior hypothalamus nucleus (AHN) region, the distribution of two receptors was dramatically different, with a higher proportion of OX1R-only cells (8.49%) compared to 1.85% in SI (Fig. 1C, D). Interestingly, much more AHN cells, 29.98% are double-positive for OX1R and OX2R, in contrast to 4.95% in the SI. The above results validated our RNAscope data, showing that approximately half of the SI cells express orexin receptors, with OX2Rs being the predominant type. This suggests the potential innervation of SI neurons by orexin.

### Local release of orexin peptide in the SI during anesthesia

After observing a rich presence of presynaptic vesicles and postsynaptic orexin receptors in the SI, we asked whether the local release of orexin participates in the anesthesia process. We employed in vivo fiber photometry using a recently developed orexin sensor - OxLight1, which was engineered by inserting a circularly permutated green fluorescent protein into the human OX2R (36). The non-Cre-dependent AAV2-OxLight1 was injected into the SI and optical fiber was implanted right above the injection site (Fig. 2A). To visualize how orexin release responds to anesthetics, we exposed the animals to 2% isoflurane, a standard dose used for rodent surgery. Upon exposure, the OxLight1 signal in the SI quickly decreased alongside behavioral arrest and stayed flat until the termination of isoflurane, after which a marked rise in the OxLight1 signal was noted, accompanied by arousal behavior (Fig. 2B). These findings demonstrated that suppression and recovery of orexin release in the SI are closely associated with anesthesia-induced behavioral changes.

**Figure 2.**
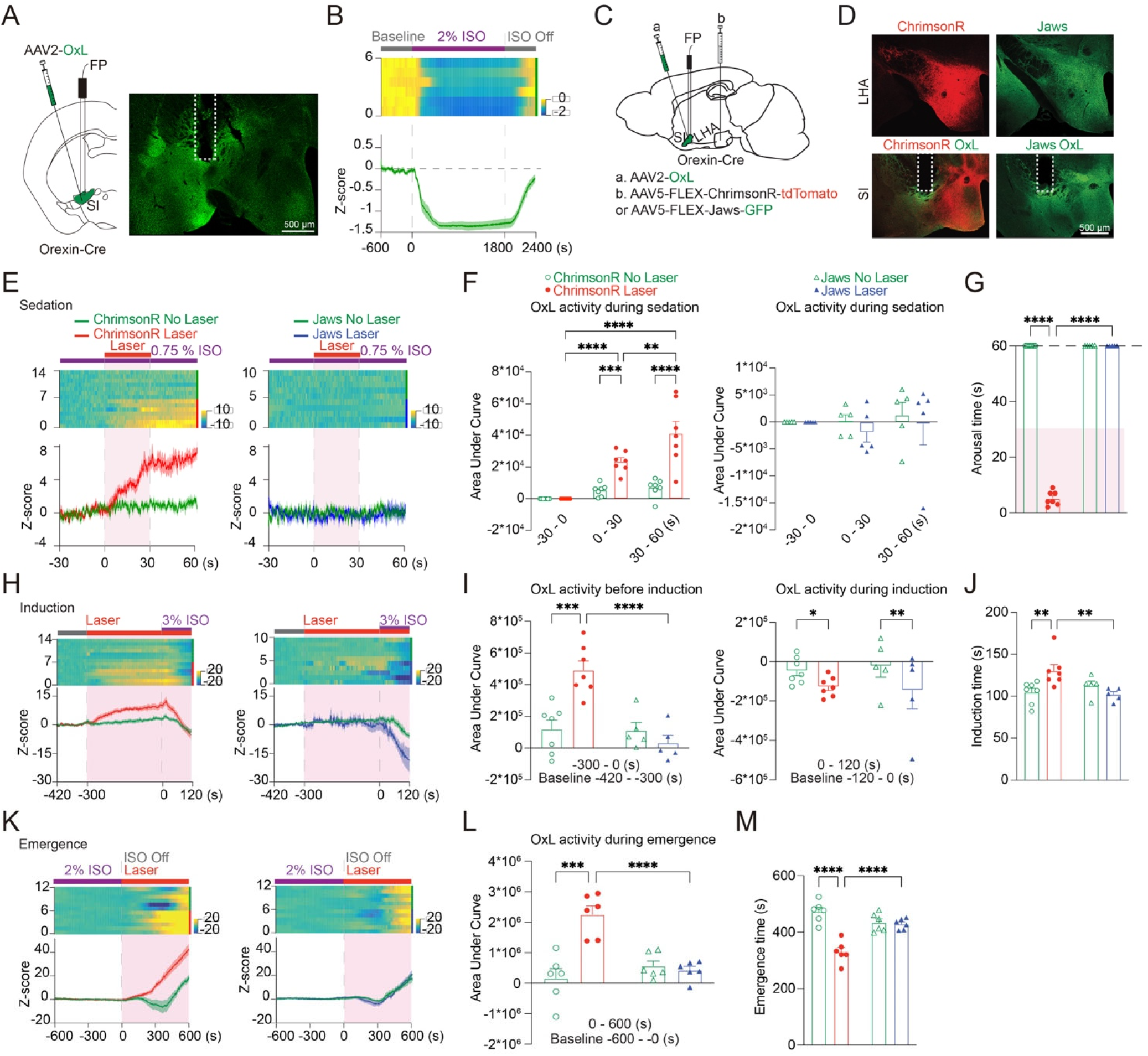
Activating LHA^OX^→SI projection increases orexin release and influences different anesthesia stages. **A**. *AAV2-OxLight1* was injected into the SI, and an optical fiber was implanted above. **B**. OxLight1 (OxL) signal changes during isoflurane anesthesia. **C, D**. Additional to *AAV2-OxLight1* injection and fiber implantation to the SI, *AAV5-FLEX-ChrimsonR-tdTomato* or *AAV5-FLEX-Jaws-GFP* was injected into the ipsilateral LHA. **E**. OxL signals during 0.75% isoflurane sedation. **F**. Area under the curve (AUC) analysis of OxL signals during sedation. ChrimsonR groups: Two-way ANOVA, the effect of laser stimulation: F_(1, 12)_ = 29.34, p = 0.0002, the effect of time: F_(2, 24)_ = 29.30, p < 0.0001, the interaction between laser stimulation and time: F_(2, 24)_ = 14.65, p < 0.0001, Tukey’s multiple comparison test, n = 7. Jaws groups: Two-way ANOVA, the effect of laser stimulation: F_(1, 8)_ = 0.3878, p = 0.55, the effect of time: F_(2, 16)_ = 0.25, p = 0.79, the interaction between laser stimulation and time: F_(2, 16)_ = 0.14, p = 0.87; Šidák’s multiple comparison test, n = 5. **G**. Arousal times from opto-stimulation during sedation. The dashed line indicates the cut-off point at 60 seconds following the laser activation. Two-way ANOVA, effect of groups: F_(1, 10)_ = 2259, p < 0.0001, effect of laser stimulation: F_(1, 10)_ = 2259, p < 0.0001, interaction between groups and laser stimulation: F_(1, 10)_ = 2259, p < 0.0001; uncorrected Fisher’s LSD analysis, ChrimsonR group, n = 7; Jaws group, n = 5. **H**. OxL signals during 3% isoflurane induction. **I**. AUC analysis of OxL signals. Pre-induction: Two-way ANOVA, the effect of groups: F_(1, 10)_ = 16.52, p = 0.0023, the effect of laser stimulation: F_(1, 10)_ = 6.58, p = 0.0281, the interaction between groups and laser stimulation: F_(1, 10)_ = 15.69, p = 0.0027; uncorrected Fisher’s LSD analysis. Post-induction: Two-way ANOVA, the effect of groups: F_(1, 10)_ = 0.003541, p = 0.9537, the effect of laser stimulation: F_(1, 10)_ = 18.83, p = 0.0015, interaction between groups and laser stimulation: F_(1, 10)_ = 0.7711, p = 0.4005; uncorrected Fisher’s LSD analysis, ChrimsonR group, n = 7; Jaws group, n = 5. **J**. Induction time comparison. Two-way ANOVA, the effect of groups: F_(1, 10)_ = 2.405, p = 0.152, the effect of laser stimulation: F_(1, 10)_ = 2.164, p = 0.172, interaction between groups and laser stimulation: F_(1, 10)_ = 12.04, p = 0.006; uncorrected Fisher’s LSD analysis, ChrimsonR group, n = 7; Jaws group, n = 5. **K**. OxL signals during 2% isoflurane emergence. **L**. AUC analysis of OxL signals during emergence. Two-way ANOVA, the effect of groups: F_(1, 10)_ = 9.168, p = 0.0127, the effect of laser stimulation: F_(1, 10)_ = 16.87, p = 0.0021, the interaction between groups and laser stimulation: F_(1, 10)_ = 21.82, p = 0.0009; uncorrected Fisher’s LSD analysis, n = 6 for both groups; **M**. Emergence times. Two-way ANOVA, the effect of groups: F_(1, 10)_ = 4.503, p = 0.0598, the effect of laser stimulation: F_(1, 10)_ = 33.95, p = 0.0002, interaction between groups and laser stimulation: F_(1, 10)_ = 29.64, p = 0.0003; uncorrected Fisher’s LSD analysis, n = 6 for both groups. *p ≤ 0.05, \*\**P* ≤ 0.01, ***p ≤ 0.001, ****p ≤ 0.0001. All data are expressed as mean ± S.E.M.

### Activating LHA^OX^→SI projection increases orexin release and reverses anesthesia sedation

To explore the relationship between orexin release, and anesthesia behavior, we combined optogenetics and fiber photometry. Specifically, we used ChrimsonR, a red light-drivable channelrhodopsin (cation channel) (37), and Jaws, a red light-drivable cruxhalorhodopsin (chloride pump) (38), to modulate orexin activity in the SI. Meanwhile, we recorded the orexin dynamics at the same location. This is achieved by injecting excitatory AAV5-FLEX-ChrimsonR-tdTomato or inhibitory AAV5-FLEX-Jaws-GFP into the LHA, as well as AAV2-OxLight1 into the SI (Fig. 2C, D). An optical fiber was implanted right above the SI, allowing us to stimulate the orexin terminals through the 635 nm channel while simultaneously recording the OxLight1 signals through the 473 nm channel.

Different types of anesthesia, from light sedation to deep general anesthesia, involve a spectrum of brain states, tailored for specific procedures to provide the best patient comfort but with minimized anesthetic exposure. We first examined orexin release during light sedation. Anesthesia was induced with 3% isoflurane for 1 minute, then switched to maintenance at 0.75% for 3 minutes followed by activation of the ChrimsonR or Jaws for 30 seconds (635 nm, 20 Hz, 20 ms) (Fig. 2E). Two-way ANOVA showed the significant effect of laser stimulation in ChrimsonR group (F_(1, 12)_ = 29.34, p = 0.0002), activation of ChrimsonR quickly increased OxLight1 signal compared to the no-laser control (0-30 s: 23426.68 ± 2502.99 vs 5336.82 ± 1424.58, Tukey’s multiple comparisons, p = 0.0007; n = 7), and the OxLight1 signal remained elevated after stimulation ended (30-60 s: 41195.01 ± 7619.86 vs 6950.38 ± 2194.26; Tukey’s multiple comparisons, p < 0.0001; n = 7; Fig. 2E, F). All animals displayed robust arousal behavior (latency to wake: 5.14 ± 0.96, Two-way ANOVA, F_(1, 10)_ = 2259, p < 0.0001, uncorrected Fisher’s LSD, p < 0.0001; n = 7) following the ChrimsonR activation (Fig. 2G). In contrast, no significant changes in OxLight1 signals were observed in the ChrimsonR no-laser control (−30-0 vs 0-30s: p = 0.469, -30-0 vs 30-60s: p = 0.285, and 0-30 vs 30-60s: p = 0.931), or the Jaws groups with or without laser stimulation (Two-way ANOVA, F_(1, 8)_ = 0.3878, p = 0.55, n = 5; Fig. 2E, F). Uncorrected Fisher’s LSD showed no significant difference in arousal behavior in the Jaws group with or without laser (p > 0.9999), the comparisons between ChrimsonR and Jaws group show no significant difference in the no-laser condition (p > 0.9999), but a significant difference in the laser condition (p < 0.0001, uncorrected Fisher’s LSD, n = 5; Fig. 2G). To verify the inhibitory function of Jaws, we tested Jaws in the acute brain slice. Whole-cell patch recordings showed that Jaws activation hyperpolarized the cell membrane potential and blocked the evoked action potentials (Supplementary Fig. 2A, B). These findings showed that activation of orexin terminals in the SI elicited the release of orexin peptides and reversed the sedation anesthesia, demonstrating the role in arousal during sedation anesthesia.

### LHA^OX^→SI activity influences the speed of anesthesia induction and emergence

General anesthesia is on the deep end of the anesthesia spectrum. When mice were exposed to isoflurane levels exceeding 2%, equivalent to general anesthesia in humans, no behavioral arousal could be triggered by optogenetic stimulation in the SI. However, the induction and emergence phases are considered the two critical and risky periods of general anesthesia. Therefore, we focused on determining how orexin activity affects these two transition stages.

We first performed anesthesia induction tests with 3% isoflurane (Fig. 2H). Recognizing the importance of time needed for orexin release to reach a plateau, we introduced a 5-minute opto-stimulation (−300-0 s) before initiating 3% isoflurane. Two-way ANOVA analysis (F _(1, 10)_ = 15.69, p = 0.0027, uncorrected Fisher’s LSD) showed that stimulating the orexin terminals via ChrimsonR in the SI caused a sustained elevation of the OxLight1 signal prior to isoflurane exposure (ChrimsonR Laser vs ChrimsonR no-laser: 490846.64 ± 58732.68 vs 119012 ± 56746.14, p = 0.0005; n = 7). In contrast, Jaws activation did not further reduce the OxLight1 signal, likely due to the low basal activity of orexin neurons (Jaws Laser vs Jaws no-laser: 31196.4 ± 49536.47 vs 110690.76 ± 51703.4, p = 0.3825; n = 5). Comparisons between ChrimsonR and Jaws show no significant difference under the no-laser condition (uncorrected Fisher’s LSD, p = 0.9192), but a significant difference under the laser condition (uncorrected Fisher’s LSD, p < 0.0001).

We then initiated 3% isoflurane induction, aligning with the clinical standard of using higher than the maintenance dose for rapid induction. The onset of 3% isoflurane (0-120 s) decreased the OxLight1 signal in both the ChrimsonR group and Jaws group (Two-way ANOVA, F_(1, 10)_ = 18.83, p = 0.0015, ChrimsonR Laser vs ChrimsonR no-laser: -127005.4 ± 19053.89 vs -45486.44 ± 25628.9, uncorrected Fisher’s LSD, p = 0.023, n = 7; Jaws Laser vs Jaws no-laser: -143743.37 ± 94474.4 vs - 20861.72 ± 57970.35, uncorrected Fisher’s LSD, p = 0.0066, n = 5) (Fig. 2H, I). Comparisons between ChrimsonR and Jaws show no significant difference in OxLight1 signal drops under the no-laser condition (uncorrected Fisher’s LSD, p = 0.7299), and the laser condition (uncorrected Fisher’s LSD, p = 0.8143; Fig. 2I). Two-way ANOVA analysis of induction times (F_(1, 10)_ = 12.04, p = 0.006) showed the ChrimsonR activation significantly prolonged the induction time, as indicated by the time to Loss of Righting Reflex (LoRR) (ChrimsonR laser vs ChrimsonR no-laser: 130.29 ± 7.21 vs 104.57 ± 5.07 s, uncorrected Fisher’s LSD, p = 0.0033, n = 7; Fig. 2J). However, no significant changes in induction time were observed in the Jaws group (Jaws laser vs Jaws no-laser: 102 ± 3.44 vs 112.4 ± 5.56 s, uncorrected Fisher’s LSD, p = 0.2201, n = 5; Fig. 2J). Comparisons between ChrimsonR and Jaws groups showed no significant difference under the no-laser condition (uncorrected Fisher’s LSD, p = 0.3626), but a significant difference under the laser condition (uncorrected Fisher’s LSD, p = 0.0031; Fig. 2J). The results suggested that activating the LHA^OX^→SI circuit slowed induction, while inhibition by Jaws had no accelerating effect.

We next performed the emergence test in the same groups of animals. The Return of the Righting Reflex (RoRR) was recorded as the emergence time. Mice were exposed to 2% isoflurane for 30 minutes, after which the isoflurane was shut off, and laser stimulation was initiated (Fig. 2K). The OxLight1 signals were completely suppressed during 2% isoflurane. Two-way ANOVA analysis of OxLight1 signals (F_(1, 10)_ = 21.82, p = 0.0009) showed opto-stimulation led to an earlier rise of OxLight1 signal during the emergence (ChrimsonR Laser vs ChrimsonR no-laser: 2,248,135 ± 282432.6 vs 160,814.28 ± 314613.39, uncorrected Fisher’s LSD, p = 0.0001; n = 6), but the activation of Jaws did not prevent the rise in the OxLight1 signal during emergence (Jaws Laser vs Jaws no-laser: 426074.15 ± 126294.89 vs 560132.37 ± 169600.85, uncorrected Fisher’s LSD, p = 0.6985; n = 6; Fig. 2K, L). Two-way ANOVA analysis of arousal behavior (F _(1, 10)_ = 29.64, p = 0.0003) showed the animals in the ChrimsonR group woke up faster with laser stimulation than the no-laser control (330 ± 15.93 vs 472.5 ± 15.31 s, uncorrected Fisher’s LSD, p < 0.0001; n = 6; Fig. 2M). In contrast, there were no significant changes in the Jaws group (Jaws Laser vs Jaws no-laser: 429 ± 7.81 vs 433.83 ± 13.23 s, uncorrected Fisher’s LSD, p = 0.7924; n =6; Fig. 2M). Comparisons between the ChrimsonR and Jaws groups showed no significant difference in the no-laser condition (uncorrected Fisher’s LSD, p = 0.0556), but a significant difference in the laser condition (uncorrected Fisher’s LSD, p < 0.0001; n = 6; Fig. 2M). Overall, the results here demonstrated that both induction and emergence were influenced by the orexin levels in the SI.

### Selective stimulation of orexin terminals activates SI neurons and induces cortical activity changes

In earlier experiments, anterograde AAV injections into the LHA orexin cell bodies allowed us to stimulate terminals in the SI. However, this approach could potentially also activate fibers passing through the SI en route to other downstream regions. To improve the specificity, we injected the retrograde AAVrg-DIO-ChR2-mCherry or AAVrg-DIO-mCherry into the SI, implanted the optic fiber targeting the same region, and attached the EEG/EMG headmount (Fig. 3A). Robust ChR2-mCherry fluorescence was observed in both the SI and LHA, confirming successful retrograde transport from orexin terminals in the SI to the LHA cell bodies (Fig. 3B).

**Figure 3.**
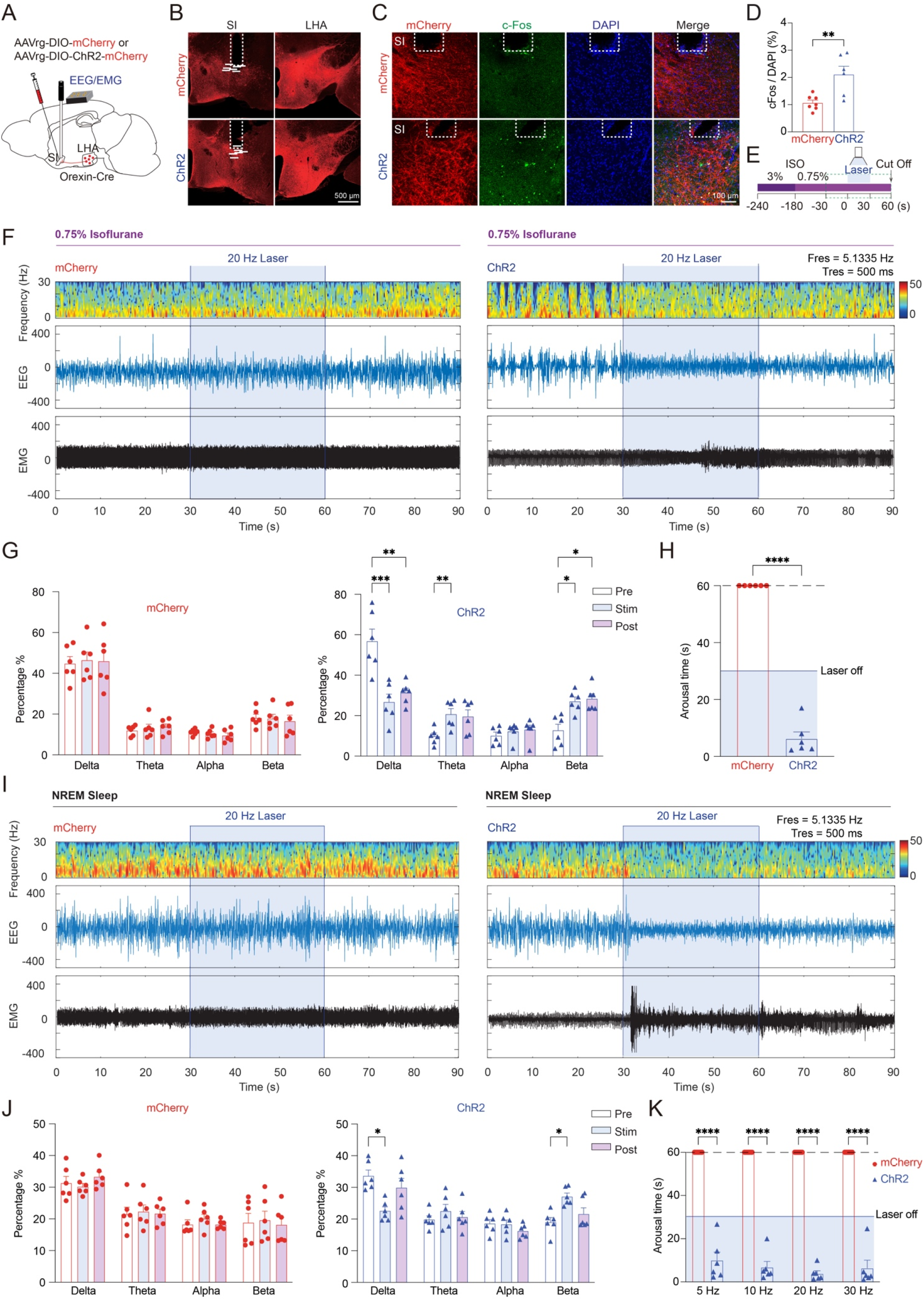
Selective activation of LHA^OX^→SI projection promotes arousal from anesthesia sedation and NREM sleep. **A**. Bilateral retrograde *AAVrg-DIO-ChR2-mCherry* or *AAVrg-DIO-mCherry* injection and optical fibers implantation into SI, with EEG/EMG headpiece attached. **B**. Expression of mCherry was detected in both the SI and LHA. Each horizontal bar below represents the location of a fiber tip. **C**. mCherry (red), cFos (green), and DAPI (blue) stainings in the SI after optogenetic stimulation. **D**. Optogenetic activation increased the cFos expression in the ChR2 group. Two-tailed unpaired *t*-test between two groups (t = 3.507, df = 11, p = 0.0049). mCherry: n = 7; ChR2: n = 6. **E**. Schematics of the arousal test under 0.75% isoflurane. **F**. EEG power spectrogram, EEG, and EMG traces, before, during, and after optogenetic activation under 0.75% isoflurane. **G**. Power percentage of EEG frequency bands. mCherry group: Two-way ANOVA, the interaction between laser stimulation and brain wave frequency: F_(6, 40)_ = 0.5352, p = 0.7782; Tukey’s multiple comparisons. ChR2 group: Two-way ANOVA, the interaction between laser stimulation and brain wave frequency: F_(6, 40)_ = 27.03, p < 0.0001); Tukey’s multiple comparisons; n = 6 for both groups. **H**. Arousal times from opto-stimulations during 0.75% isoflurane. ChR2 group: 6.33 ± 2.3 s. mCherry group: no arousal. Two-tailed unpaired *t*-test (t = 23.32, df = 10, p < 0.0001), n = 6 for both groups. **I**. EEG power spectrogram, EEG, and EMG traces before, during, and after optogenetic activation under NREM sleep. **J**. Power percentage of EEG frequency bands. mCherry group: Two-way ANOVA, the interaction between laser stimulation and brain wave frequency: F_(6, 48)_ = 0.6133, p = 0.7182; Tukey’s multiple comparisons; n = 6; ChR2 group: Two-way ANOVA, the interaction between laser stimulation and brain wave frequency: F_(6, 40)_ = 8.26, p < 0.0001, Tukey’s multiple comparisons; n = 6. **K**. Arousal times from NREM sleep responding to different frequencies of opto-stimulations. Two-way ANOVA, the effect of groups: F_(1, 10)_ = 491, p < 0.0001); Bonferroni’s multiple comparisons, n = 6 for both groups. *p ≤ 0.05, \*\**P* ≤ 0.01, ***p ≤ 0.001, ****p ≤ 0.0001, All data are expressed as mean ± S.E.M.

To confirm the optogenetic activation of SI neurons, we used cFos staining, a common marker of recent neuronal activity. 470 nm laser stimulations (20 Hz, 20 ms, 1 s on, 1 s off, 30 minutes) indeed induced a significant increase of cFos staining in the SI compared to the control animals expressing only mCherry (ChR2: 2.11 ± 0.30, n=6; mCherry: 1.07 ± 0.10, n=7; two-tailed unpaired t-test, t = 3.507, df = 11, p = 0.0049; Fig. 3C, D), indicating that selective stimulation activation of orexin terminals effectively activated the SI neurons.

EEG recordings, which are widely used to monitor anesthesia depth, were taken to determine the effect of LHA^OX^→SI circuit activation on cortical activity. We exposed the mice expressing ChR2-mCherry or mCherry to 0.75% isoflurane anesthesia and performed the arousal tests (Fig. 3E). Similar to the ChrimsonR activation, the 473 nm stimulation of ChR2 (20 Hz, 20 ms, 30 s) reliably awakened the mice from 0.75% isoflurane within 6.33 ± 2.3 seconds after stimulation onset (two-tailed unpaired t-test, t = 23.32, df = 10, p < 0.0001, n = 6; Fig. 3H). EEG from the ChR2 group showed a transition to higher frequency and lower amplitude waveforms, marked by a decrease in delta (pre: 56.86 ± 6.00, stim: 26.78 ± 3.88, post: 31.37 ± 2.24; Tukey’s multiple comparisons, pre vs stim, p = 0.0006, pre vs post, p = 0.0049; n = 6) and increases in theta (pre: 9.81 ± 1.61, stim: 20.76 ± 2.74, post: 19.6 ± 3.29; Tukey’s multiple comparisons, pre vs stim, p = 0.0021, pre vs post, p = 0.0563; n = 6) and beta (pre: 12.76 ± 3.04, stim: 27.13 ± 2.12, post: 28.4 ± 2.45; Tukey’s multiple comparisons, pre vs stim, p = 0.0125, pre vs post, p = 0.0342; n = 6), consistent with arousal behavior (Two-way ANOVA, F_(6, 40)_ = 0.5352, p = 0.7782 for mCherry group, F_(6, 40)_ = 27.03, p < 0.0001 for ChR2 group; Fig. 3F, G).

### Selective activation of LHA^OX^→SI projection promotes wakefulness from NREM sleep

Anesthesia and sleep share similar neural circuits. We then asked whether the LHA^OX^→SI circuit regulates sleep/wake control. We stimulated free-moving animals with 470 nm laser pulses (20 Hz, 20 ms, 30 seconds) after they entered NREM sleep, as determined by their EEG waveform. Consistently, the EEG spectrum in the ChR2 group showed a decrease in the delta band (pre: 33.62 ± 1.83, stim: 22.55 ± 1.27, post: 29.89 ± 2.93; Tukey’s multiple comparisons, pre vs stim, p = 0.0113, pre vs post, p = 0.2002; n = 6) but an increase in beta (pre: 19.04 ± 1.44, stim: 27.1 ± 1.14, post: 21.59 ± 1.97; Tukey’s multiple comparisons, pre vs stim, p = 0.0422, pre vs post, p = 0.2851; n = 6), indicating arousal from NREM sleep (Two-way ANOVA, F_(6, 40)_ = 8.26, p < 0.0001, Fig. 3I, J). In contrast, the mCherry control mice showed no response to the optogenetic stimulation (Two-way ANOVA, F_(6, 48)_ = 0.6133, p = 0.7182; n = 6; Fig. 3I, J). We tested four laser frequencies (5 Hz, 10 Hz, 20 Hz, and 30 Hz), all of which induced rapid arousal. While lower-frequency simulations produced slightly slower responses (5 Hz: 10 ± 3.79 s, n = 6) compared to higher frequencies (10 Hz: 6.78 ± 2.7 s, 20 Hz: 3.83 ± 1.32 s, 30 Hz: 6.35 ± 3.70 s, n = 6), they were statistically insignificant (Two-way ANOVA, F_(1, 10)_ = 491, p < 0.0001, Bonferroni’s multiple comparisons, p < 0.0001, Fig. 3K). These findings highlighted that the activation of orexin terminals in the SI induces arousal from NREM sleep, similar to their role in promoting arousal from anesthesia.

### Selective activation of LHA^OX^→SI projection induces analgesia

Analgesia is an integral part of anesthesia. As demonstrated by previous studies that activating orexin neurons in LHA can induce analgesia (39), we asked whether SI is involved. A 55°C hotplate test was used to assess the sensitivity to thermal pain. Mice expressing ChR2-mCherry or mCherry were placed onto the hotplate after 5 minutes of continuous opto-stimulation (473 nm, 20 ms, 20 Hz, 1 s on, 1 s off) and the withdrawal latency (time to paw withdrawal) was recorded (Fig. 4A). With laser stimulation, ChR2 mice showed a significantly longer withdrawal latency (11.48 ± 0.95 s, Two-way ANOVA, F_(1, 10)_ = 13.19, p = 0.0046, n = 6) than the same animals without laser stimulation (8.62 ± 0.39 s, uncorrected Fisher’s LSD, p = 0.0074), and the mCherry group with laser stimulation (7.68 ± 0.79 s, uncorrected Fisher’s LSD, p = 0.001; n = 6). No significant difference was observed in the mCherry group with or without laser stimulation (uncorrected Fisher’s LSD, p = 0.1037; Fig. 4B).

**Figure 4.**
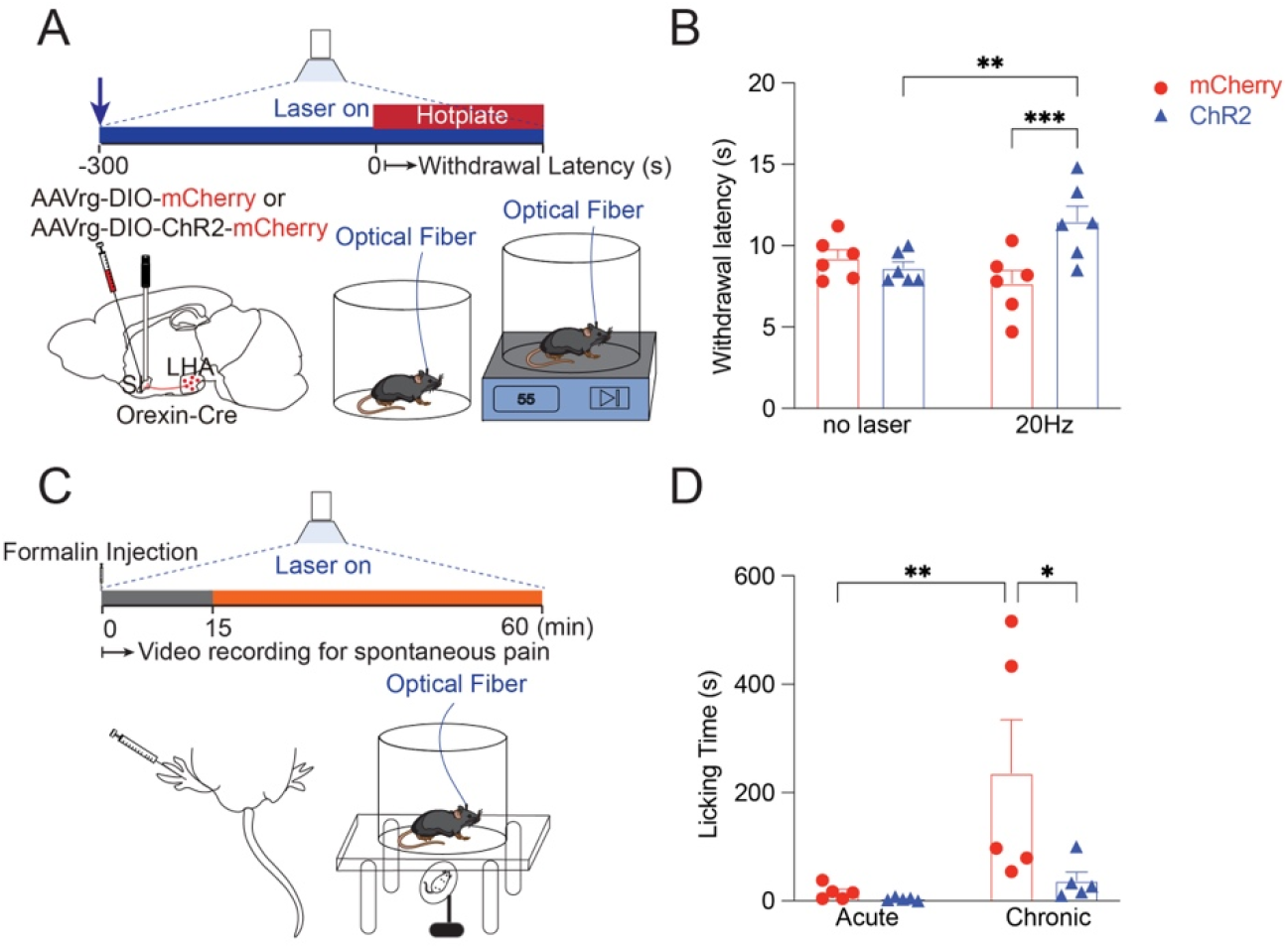
Selective activation of LHA^OX^→SI projection induces analgesia. **A**. Retrograde *AAVrg-DIO-ChR2-mCherry* or *AAVrg-DIO-mCherry* was injected into the SI of *Orexin-Cre* mice bilaterally and optical fibers were implanted above. The mice were tested on a 55°C hotplate after 5 minutes of opto-stimulation with a 473 nm laser. **B**. Optogenetic activation of LHA^OX^→SI increased the withdrawal latency to the thermal stimulation for the ChR2 laser group (20Hz laser vs no-laser: 11.48 ± 0.95 vs 8.62 ± 0.39 s, uncorrected Fisher’s LSD, p = 0.0074), but not the mCherry groups (20Hz laser vs no-laser: 7.68 ± 0.79 vs 9.22 ± 0.53 s, uncorrected Fisher’s LSD, p = 0.1037). Two-way ANOVA, the interaction between laser stimulation and group F_(1, 10)_ = 13.19, p = 0.0046; uncorrected Fisher’s LSD analysis of mCherry and ChR2 in 20Hz laser showed a significant difference (p = 0.001); n = 6 for both groups. **C**. Formalin test. The left hind paws of the mice were injected with 10 ul 5% formalin followed by a 60-minute video recording with continuous opto-stimulation. **D**. Optogenetic activation of LHA^OX^→SI reduced the licking time for the ChR2 group during the chronic phase (mCherry vs ChR2: 235.8 ± 98.57 vs 37.2 ± 16.21 s, uncorrected Fisher’s LSD, p = 0.0127), but not the acute phase (mCherry vs ChR2: 15.6 ± 6.22 vs 3.6 ± 1.36 s, uncorrected Fisher’s LSD, p = 0.8675). Two-way ANOVA, the interaction between laser stimulation and group F_(1, 16)_ = 3.476, p = 0.0807, uncorrected Fisher’s LSD analysis of different phases in the mCherry group showed a significant difference (p = 0.0067), but no difference in ChR2 group (p = 0.6414); n = 5 for both groups. *p ≤ 0.05, **p ≤ 0.01, ***p ≤ 0.001. All data are expressed as mean ± S.E.M.

We also performed a formalin test to evaluate the sensitivity to inflammatory pain. 10 μl 5% formalin was injected subcutaneously into the dorsal part of the hind paw followed by immediate laser stimulation, and the pain behaviors (licking) were recorded for 1 hour (Fig. 4C). The first 15-minute phase, considered an acute phase, likely reflects direct nerve terminal stimulation, whereas the following 45 minutes, considered as the chronic phase, reflects the inflammatory pain responses. Although no significant difference was observed during the acute phase (mCherry vs ChR2: 15.6 ± 6.22 vs 3.6 ± 1.36 s, Two-way ANOVA, F_(1, 16)_ = 3.476, p = 0.0807, uncorrected Fisher’s LSD, p = 0.8675; n = 5), activation of orexin terminals in the SI dramatically reduced the licking time of the ChR2 mice during the chronic phase (mCherry vs ChR2: 235.8 ± 98.57 vs 37.2 ± 16.21, uncorrected Fisher’s LSD, p = 0.0127; n = 5; Fig. 4D). These results suggested that the LHA^OX^→SI projection is indeed involved in analgesia in thermal and inflammatory pain.

### Both EPSCs and IPSCs are elicited by stimulating orexin terminals in the SI

After we studied the functional roles of the LHA^OX^→SI circuit, we asked about the synaptic properties of orexin innervations in the SI. AAV5-FLEX-ChrimsonR-tdTomato was injected into the LHA, and the brain slices containing the SI region were used for whole-cell patch-clamp recordings (Fig. 5A, B). Upon applying a short train of the red laser pulses (635 nm, 20 Hz, 20 ms), both excitatory (EPSC) and inhibitory postsynaptic currents (IPSC) were detected in the SI neurons (Fig 5C).

**Figure 5.**
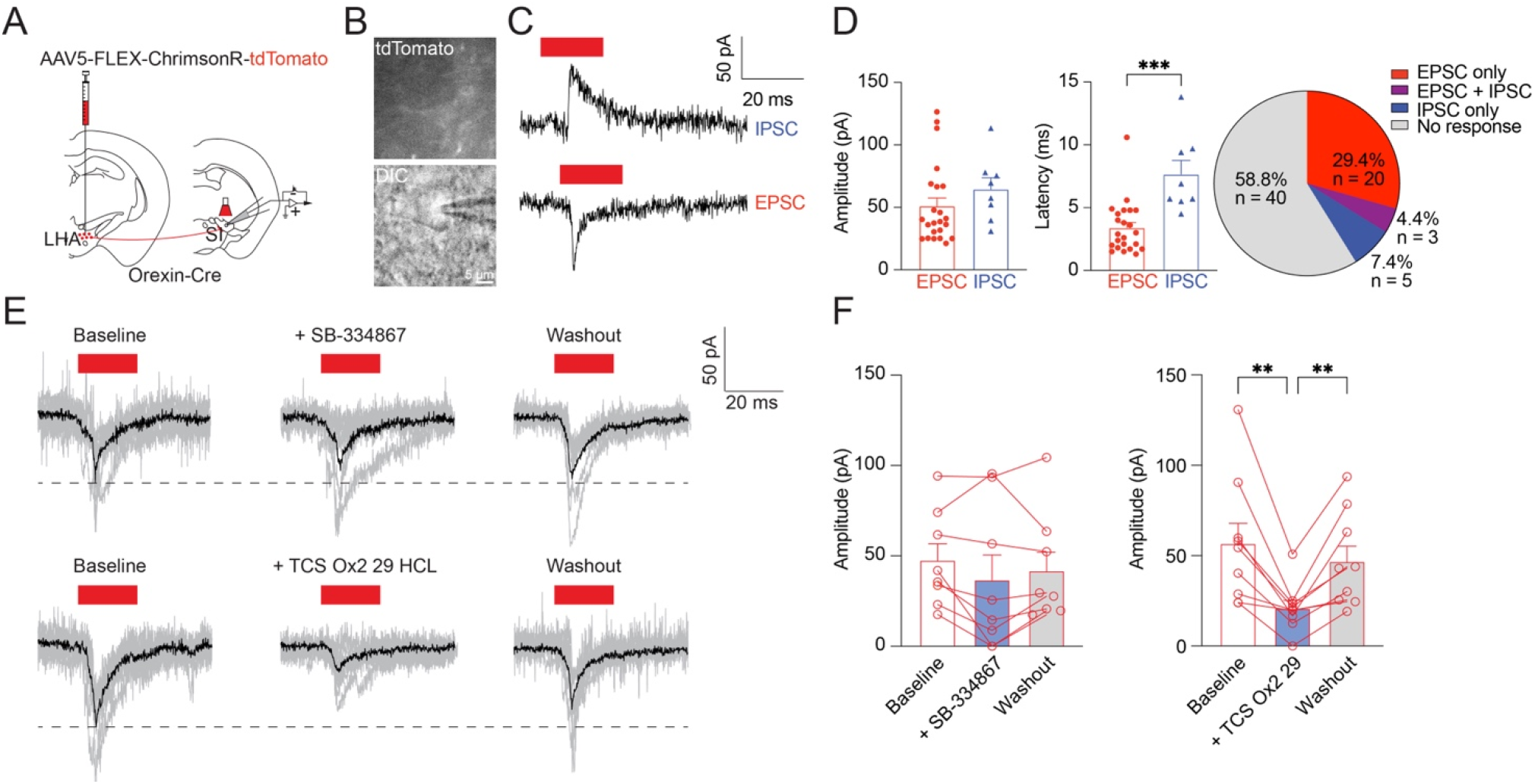
SI neurons receive monosynaptic excitatory inputs from orexin neurons. **A**. *AAV5-FLEX-ChrimsonR-tdTomato* was injected into the LHA of *Orexin-Cre* mice, and patch-clamp whole-cell recordings were performed on the SI neurons. **B**. Abundant tdTomato-positive fibers surrounding the SI neurons. **C**. Both EPSCs and IPSCs were elicited by photostimulation (635 nm). **D**. Amplitude (EPSC: 51.03 ± 6.58 pA, n = 23; IPSC: 64.54 ± 9.12 pA, n = 8; Two-tailed unpaired *t*-test, t = 1.087, df = 29, p = 0.2858), latency (EPSC: 3.39 ± 0.43 ms, n = 23; IPSC: 7.65 ± 1.1 ms, n = 8; Two-tailed unpaired *t*-test, t = 4.393, df = 29, p = 0.0001), and distribution. **E, F**. Laser-evoked EPSCs were blocked by OX2R antagonist - TCS OX2 29, but not as much by OX1R antagonist - SB-334867. One-way ANOVA, the effect of SB-334867, F_(1.536, 10.75)_ = 1.944, p = 0.1928, Tukey’s multiple comparisons of the amplitude of the baseline (47.71 ± 9.39 pA), SB-334867 (36.81 ± 14.12 pA), and washout (41.92 ± 10.68 pA) showed no significant difference (baseline vs SB-334867: p = 0.2712, SB-334867 vs washout: p = 0.6891); n = 8. One-way ANOVA analysis of the effect of the TCS OX2 29, F_(1.262, 10.09)_ = 17.38, p = 0.0012, Tukey’s multiple comparisons of baseline (56.76 ± 11.69 pA), TCS OX2 29 (21.1 ± 4.48 pA), and washout (46.88 ± 8.71 pA) showed TCS OX2 29 significantly decreased the amplitude of the laser evoked EPSCs (baseline vs TCS OX2 29: p = 0.006, TCS OX2 vs washout: p = 0.0058); n = 9. N stands for cell numbers, each data point represents an average of five traces from a single cell. **p ≤ 0.01, ***p ≤ 0.001. All data are expressed as mean ± S.E.M.

Among all tested neurons, 41.2% responded to the laser stimulation. Most responses were EPSCs (29.4%, n = 20), with fewer IPSCs (7.4%, n = 5) and both EPSCs and IPSCs (4.4%, n = 3). The amplitudes of EPSCs and IPSCs were similar, around 50 pA. The EPSCs had shorter latencies of less than 5 ms (3.39 ± 0.43 ms, n = 23) indicating they were monosynaptic connections (40). While the IPSCs had a significantly delayed response (7.65 ± 1.1 ms, two-tailed unpaired t-test, t = 4.393, df = 29, p = 0.0001, n = 8, Fig 5D), suggesting an indirect innervation by the orexin neurons. Furthermore, the EPSCs can be reversibly blocked by OX2R antagonist - TCS-OX2-29 (One-way ANOVA, F_(1.262, 10.09)_ = 17.38, p = 0.0012, Tukey’s multiple comparisons, baseline vs TCS OX2 29: p = 0.006, TCS OX2 vs washout: p = 0.0058; n = 9), but not OX1R antagonist - SB-334867 (One-way ANOVA, F_(1.536, 10.75)_ = 1.944, p = 0.1928, Tukey’s multiple comparisons, baseline vs SB-334867: p = 0.2712, SB-334867 vs washout: p = 0.6891; n = 8; Fig 5E, F). This indicated that OX2R but not OX1R in the SI are involved in the excitatory synaptic transmission. The IPSCs exhibited similar responses to the blockers (Supplementary Fig. 2C, D).

### Downstream neurons of the LHA^ox^ in the SI

We further investigated the downstream neuronal types in the SI that are innervated by orexin terminals. Using RNAscope, four neuronal types marked by different mRNAs - ChAT (Cholinergic), Slc17a6 (vGlut2, glutamatergic), Slc32a1 (vGAT, GABAergic), and TH (Catecholaminergic), were examined for colocalization with OX2R mRNA.

Although SI has been known as the major source of cholinergic neurons, only a small portion of SI cells were cholinergic neurons (approximately 4.13% of all SI cells, average of ChAT stainings from Fig. 6A-C). Meanwhile, there were significantly more glutamatergic neurons (11.96%; Fig. 6A, D) and GABAergic neurons (28.17%; Fig. 6B, E), along with a smaller population number of catecholaminergic neurons in the SI (1.66%; Fig. 6C, F). Due to the lack of capability to co-stain four probes all at once, we used the ChAT probe as a reference in all sets of RNAscope staining to compare the ratios between different neuronal types. The analysis revealed ratios of approximately 2.81 for vGlut2/ChAT, 8.86 for vGAT/ChAT, and 0.73 for TH/ChAT (Supplementary Fig. 3A, B).

**Figure 6.**
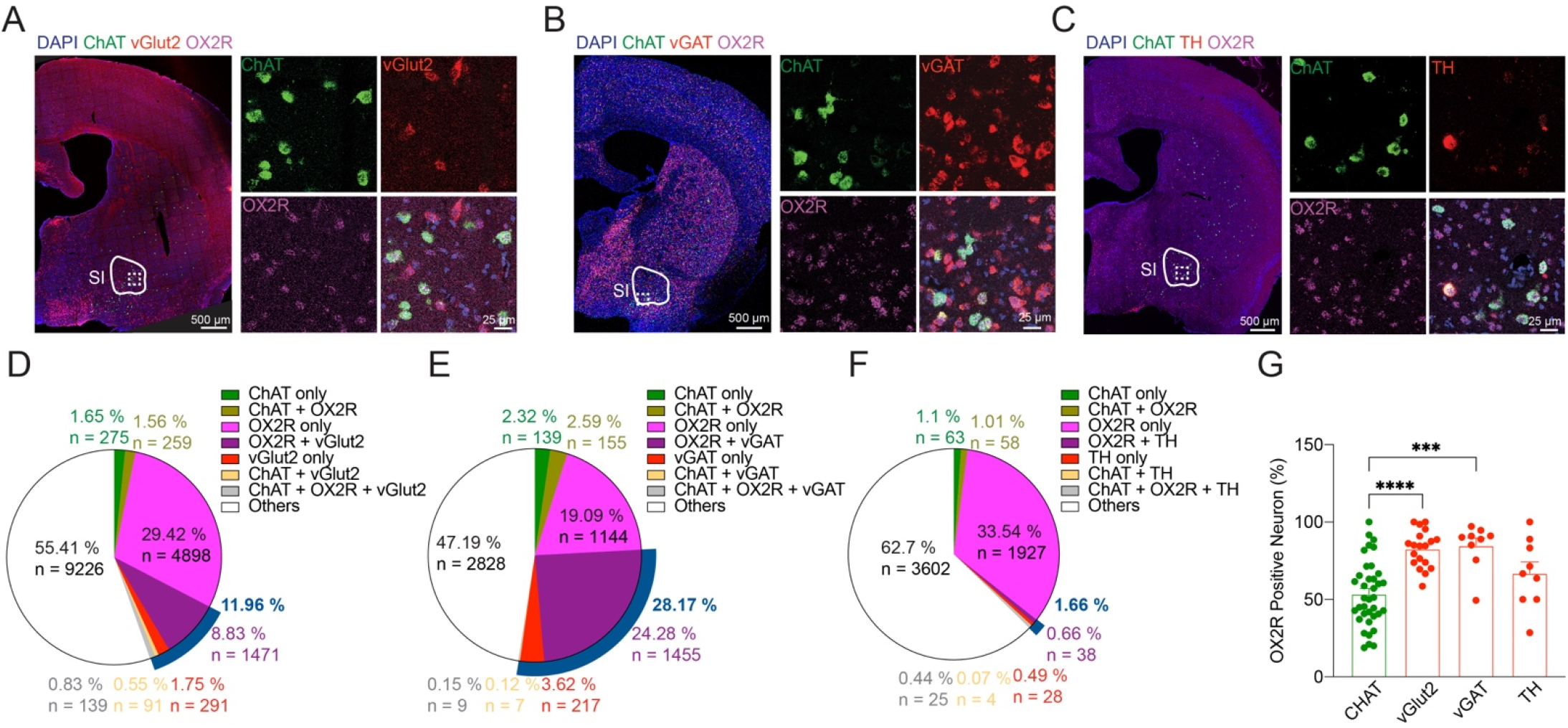
The downstream neurons of the LHA^OX^ in the SI. **A-C**. RNAscope fluorescence for different types of neurons - ChAT, vGlut2, vGAT, TH and their stainings with OX2R in SI. **D-F**. Colocalizations of ChAT, OX2R, and vGlut2 (D. 19 slices from 5 brains); or vGAT (E. 9 slices from 5 brains), or TH (F. 9 slices from 5 brains). **G**. OX2R distributions in ChAT, vGlut2, vGAT, and TH neurons in SI. One-way ANOVA, F_(3, 70)_ = 13.66, p < 0.001, Tukey’s multiple comparisons test for OX2R+ChAT/ChAT (53.6 ± 3.46 %), OX2R+vGlut2/vGlut2 (82.64 ± 2.73 %), OX2R+vGAT/vGAT (84.8 ± 4.88 %), OX2R+TH/TH (66.95 ± 7.38 %) showed OX2R are distributed more in vGlut2 (p < 0.0001) and vGAT (p = 0.0001) than ChAT neurons. ChAT neurons were counted from 37 brain slices of 5 mice, vGlut2 neurons were from 19 brain slices of 5 mice, and vGAT and TH neurons were from 9 brain slices of 5 mice. ***p ≤ 0.001, ****p < 0.0001. All data are expressed as mean ± S.E.M.

Furthermore, we calculated the percentage of OX2R colocalization in each one of these neuronal types. Among ChAT neurons, 53.6% were double positive for OX2R. Similarly, 82.64% of vGlut2 neurons, 84.8% of vGAT neurons, and 66.95% of TH cells co-expressed OX2R (Fig. 6G). On the other hand, among the OX2R+ neurons, 22.34% were double positive for vGlut2, 51.38% for vGAT, 2.47% for TH, and 4.53% for ChAT (Supplementary Fig. 3C, D). Taken together, these data indicated that all four neuronal types in the SI expressed OX2R, with vGAT neurons being the most abundant and showing the highest level of OX2R expression.

## Discussion

To study the role of the orexin circuit in anesthesia and analgesia, we investigated the LHA^OX^→SI circuit in Orexin-Cre mice with cutting-edge technologies such as optogenetics, in vivo fiber photometry, EEG/EMG, RNAscope, and electrophysiology tools. Our findings showed that activation of LHA^OX^→SI projections effectively awaken animals from both NREM sleep and isoflurane anesthesia, delays anesthesia induction, accelerates recovery, and enhances tolerance to thermal and formalin-induced inflammatory pain. These effects are primarily mediated by OX2Rs, which are expressed on ChAT, vGlut2, and vGAT neurons in the SI.

The arousal system consists of multiple neural pathways that safeguard survival. On one end of the arousal spectrum is sleep, while hypervigilance lies at the opposite extreme, with varying degrees of alertness in between. Anesthesia similarly involves different levels of sedation depending on the procedure. The orexin circuit is a key modulator in arousal control through its extensive projections in the brain. The SI is part of the basal forebrain (BF), located between the amygdala and the globus pallidus, and is known as the major source of cholinergic inputs to the cortex, relaying ascending arousal projections from the brainstem to the cortex (41). Other arousal-related systems include the adrenergic neurons in the LC, the histaminergic neurons in the TMN, the serotonin neurons in the raphe nuclei, and the dopaminergic neurons in the VTA, all of which have intricate connections with orexin neurons (42).

When one arousal system is not working properly, other systems may compensate to maintain the appropriate wakefulness. This may explain why anesthesia induction and emergence speed were not significantly altered when Jaws was activated in the SI. On the contrary, when one of the arousal systems, orexin in this case, was activated, the system arousal level was elevated therefore the induction of anesthesia was delayed, emergence was accelerated, and reanimation can be triggered during light anesthesia. The dynamics of OxLight1 signals further supported the association between increased orexin release and changes in arousal during anesthesia.

The SI contains a diverse population of neurons, including cholinergic, GABAergic, and glutamatergic neurons. Our RNAscope data showed that around 28.17% of SI cells are vGAT neurons, followed by 11.96% vGlut2 and 4.13% ChAT. Although the discussion about catecholaminergic neurons was not usually associated with BF (43), we did identify 1.66% TH neurons in the SI. Our data suggested a greater abundance of GABAergic and Glutamatergic neurons compared to cholinergic neurons in the SI. Future studies should focus on selectively activating these different types of neurons in the SI to further explore their roles in arousal and analgesia.

Approximately half of SI cells expressed the orexin receptors, as shown by RNAscope data, with optogenetic stimulation of orexin terminals eliciting mostly EPSCs (29.4%) and some IPSCs (7.4%). The short latency and good signal-to-noise ratio of EPSCs indicate direct monosynaptic connections, with OX2R, but not OX1R receptors involved. The longer latency of IPSCs suggests indirect connections, through inhibitory interneurons. Previous studies have shown that orexin neurons co-release glutamate and dynorphin in addition to the orexin peptide, which may explain why the EPSCs cannot be fully blocked by OX2R antagonist.

Pharmacological studies have reported diverse distributions of orexin receptors (44). Our data showed that OX2R is the dominant receptor in the SI, with 96% of orexin receptor-positive cells expressing OX2R. A survey of other sites in our brain slices showed that the LC primarily contains OX1R, whereas the AHN and PVT have both OX1R and OX2R present (Fig. 1C, Supplementary Fig. 1). It’s well-known that the LC is the main source of noradrenaline in the brain, and the SI is the main source of acetylcholine. Although both are important in arousal regulation, likely influencing different aspects of arousal, they achieve their functions through different orexin receptors. The OX1R in the LC has been shown to be involved in fear memory consolidation (45) and reward-seeking behavior (46). Therefore, the functional differences between OX1R and OX2R merit further investigation.

Out of all OX2R+ neurons in the SI, 51.38% are vGAT GABAergic neurons, 22.34% are vGlut2 glutamatergic neurons, 4.53% are ChAT cholinergic neurons, and 2.47% are TH catecholaminergic neurons (Supplementary Fig. 3C, D). GABAergic neurons have complex heterogeneous subgroups that exhibit tonic or phasic activity, which could be associated with gating activity in cortical neurons. 40% of GABAergic neurons in BF increased their discharge rate with somatosensory evoked cortical activation, while the rest became halted (47). The BF GABAergic neurons are typically smaller than Cholinergic neurons, and function not only as inhibitory interneurons to regulate nearby neurons but also as projecting neurons to the neocortex (48). Somatostatin+ GABAergic neurons promote sleep, while the parvalbumin+ GABAergic and glutamatergic neurons promote arousal (49). The fast-spiking parvalbumin+ neurons have been recognized as important players in generating gamma oscillation (50).

Cholinergic, glutamatergic, and parvalbumin+ GABAergic neurons in the BF are highly active during wakefulness and REM sleep, and optogenetic activation of each type can lead to sleep-wake transition (32). BF cholinergic neurons are well known for their role in arousal (51,52). Another study showed that BF glutamatergic neurons regulate sleep homeostasis. Ablation of vGlut2 neurons in BF by AAV-DIO-Caspase-3 caused diminished adenosine accumulation during wake and REM sleep, resulting in the animals’ increased wakefulness at night (53). Reciprocal innervation of LHA^OX^ by the BF GABAergic and glutamatergic neurons, but not by cholinergic neurons has been shown (54). Future studies are needed to clarify the roles of different SI neuronal types in arousal control.

Orexin activation, through chemogenetics or optogenetics, has been shown to modulate pain perception (20,25,55). The orexin system targets various brain regions involved in pain processing, including the periaqueductal gray (56,57), rostral ventromedial medulla (23), medial preoptic area (58), and ventral tegmental area (59). Here, we demonstrate that SI is also involved in pain control. The pain pathway involves peripheral reception, spinal cord ascending transmission, and brain interpretation. Orexin neurons project widely throughout the brain and spinal cord, suggesting that multiple sites may be involved in processing pain signals. More research is needed to reveal the mechanisms by which the LHA^ox^→SI pathway regulates pain.

## Materials and Methods

### Animals

115 mice used in this experiment were 6-12 weeks of age, and weight 18-32g. The Orexin-Cre-2A-EGFP line was a gift from Dr. Akihiro Yamanaka, Nagoya University, Japan (8), and was bred onto the C57BL/6 genetic background. The wild-type C57/BL6 mice were purchased from Jackson Laboratories (stock No: 000664). 45 Orexin-Cre mice were used in the experiments for NREM sleep, arousal, and emergence behavior tests with optogenetic stimulation and EEG/EMG recording. 26 Orexin-Cre mice were used in the experiments for investigating the orexin release during anesthesia sedation, induction, and emergence procedure with optogenetic stimulation and fiber photometry recording. 30 Orexin-Cre mice were used for the hot plate and formalin tests. 9 Orexin-Cre mice were used for the brain slice recording. 5 wild-type mice were used for the RNAscope. All experimental procedures were approved by the Institutional Animal Care and Use Committee, University of California, San Francisco. Mice were maintained in a strictly controlled environment with ad libitum access to food and water. The light cycle starts at 7:00 AM and ends at 7:00 PM, the dark cycle runs from 7:00 PM to 7:00 AM. The temperature was controlled between 20 °C to 22 °C. 10 Orexin-Cre mice were used in the experiments for neural tracing.

### Virus Injection

AAV1-phSyn1(S)-FLEX-tdTomato-T2A-SypEGFP-WPRE (51509-AAV1), AAVrg-hSyn-DIO-mCherry (50459-AAVrg), AAV-EF1a-double floxed-hChR2(H134R)-mCherry-WPRE-HGHpA (20297-AAVretro), AAV-Syn-DIO-ChrimsonR-tdTomato (62723-AAV5), and AAV5-CAG-FLEX-rc [Jaws-KGC-GFP-ER2] (84445-AAV5) were purchased from Addgene. The plasmid of AAV2-Oxlight1 (169792; Addgene) was purchased from Addgene, and the virus was homemade according to the published protocol from the Salk Institute (https://www.salk.edu/science/core-facilities/viral-vector-core/services/). All AAV viruses were aliquoted and stored at −80 °C. Each virus aliquot will not be re-frozen after each thaw.

Intracranial injections were performed under balanced anesthesia using a robot stereotaxic instrument (Neurostar, Germany). Meloxicam (5 mg/kg, s.c. injection), ampicillin (10-20 mg/kg, s.c. injection), buprenorphine (0.1 mg/kg, s.c. injection), 0.25% bupivacaine (0.025 ml, s.c. injection) and isoflurane (Henry Schein Animal Health) were given during the procedure. A 2 μl Hamilton™ Neuros™ 7000 Series Modified Microliter Syringes with Knurled Hub Needle (14-815-903; Hamilton Company) was used for the injections, and 0.5 μl virus was delivered over 10 minutes. The needle was left in situ for 5 minutes and slowly withdrawn. The skin was stapled after injection surgery using the BD Autoclip™ Wound Closing System (22-275998; Fisher Scientific).

The injection coordinates were as follows: lateral hypothalamic area (AP, -1.46 mm, ML, ± 0.95 mm, and DV, 5.1-5.0 mm), substantia innominata (AP, -0.6 mm, ML, ± 1.75 mm, and DV, 4.8-4.7 mm).

### Fiber Implantation

For optogenetic and EEG experiments, all mice were implanted with bilateral optical fibers (400 μm Core, 0.39 N.A.; R-FOC-BL400C-39NA; RWD Life Science) into the substantia innominata (AP, - 0.6 mm, ML, ± 1.75 mm, and DV, 4.7 mm).

For fiber photometry experiments, all mice were implanted with unilateral optical fiber (200 μm Core, 0.39 N.A.; R-FOC-BL200C-39NA; RWD Life Science) into the left side of the substantia innominata (AP, -0.6 mm, ML, - 1.75 mm, and DV, 4.7 mm).

### EEG/EMG Implantation

After receiving bilateral stereotaxic viral injections and optical fiber implantation, the animals received EEG/EMG device headmounts (8201, Pinnacle Technologies) implantation during the same procedure. A 23-gauge surgical needle was used to drill four guide holes, including two frontal cortical areas (AP, 1 mm, ML, ± 1.25 mm) and two parietal areas (AP, -3 mm, ML, ± 2.5 mm). Four screws with wires (8493, Pinnacle Technologies) were placed into the skull through the holes. The wires were then soldered onto a six-pin connector EEG/EMG headmount. EMG leads were placed into the neck muscle. The headmount was secured with dental cement (C&B Metabond® Quick Adhesive Cement System, Parkell) onto the skull. Behavioral experiments were conducted 4 weeks after surgery to allow for sufficient recovery.

### EEG recording with optogenetic manipulation

Mice were anesthetized with 3% isoflurane for 1 minute in an induction chamber and then attached to the EEG/EMG system (8200-K1-SL 2 EEG/1 EMG Mouse System for Sleep, Pinnacle Technologies), and the bilateral optical fibers through the black ceramic ferrule mating sleeves (R-MS-1.25, RWD Life Science Inc). The bifurcated fiber bundle (Ø400 μm Core, 0.39 NA, FC/PC to Ø1.25 mm Ferrules, 1 m Long; BFYL4LF01; Thorlabs) was connected to a 473nm laser stimulator (Aurora-200, Newdoon). The mice were acclimated for 24 hours before EEG-related behavioral testing, and during each testing session, they were given another 30 minutes of acclimation before data collection.

### Fiber photometry recording with optogenetic manipulation

Mice were anesthetized with 3% isoflurane for 1 minute in an induction chamber and the low-autofluorescence mono-fiber-optic patch cord (MFP_400/440/3000-0.37_1m_FCM-MF1.25(F)_LAF, Doric Lenses Inc.) was connected to the R821 tricolor multichannel fiber photometry system (RWD Life Science Inc.). The green fluorescence of calcium signals was evoked by 470 nm. 410 nm was used to acquire reference signals to eliminate noise. A 635 nm red laser stimulator (Intelligent Optogenetics System IOS-635, RWD Life Science Inc.) was connected to the R821 fiber photometry system to manipulate the neuronal activities of mice expressing ChrimsonR opsin and record the calcium activities simultaneously. The mice were given more than 30 minutes of acclimation before data collection. The parameters of optogenetic activation are described below in the behavioral tests.

### Immunohistochemistry

The mice were transcardially perfused 4-8 weeks after virus injection with cold PBS followed by a cold solution of 4% paraformaldehyde (PFA, Electron Microscopy Services) in PBS. Brains were removed and post-fixed in 4% PFA for 6 h at 4 °C before being transferred to 30% sucrose for at least 2 days of dehydration until the tissue stayed afloat. A series of 35 μm coronal/sagittal brain slices were sliced using a cryostat (Leica CM3050S).

For immunohistochemistry staining, brain slices were blocked with blocking solution (5% donkey serum, 3% BSA, and 0.3% Triton-X100 in PBS) for 1 hour at room temperature, followed by incubation with primary antibodies at 4°C for 16-24 hours. After the brain slices were washed three times with PBS (10 minutes each time), the slices were incubated with secondary antibodies for 2 hours at room temperature followed by washing 3 times. Finally, the slices were covered with DAPI Fluoromount-G mounting medium (0100-20, SouthernBiotech).

The antibodies were diluted in a blocking solution as follows. mCherry: Primary, chicken anti-mCherry (1:400, NBP2-25158, Novus Biologicals). Secondary, Cy3-conjugated AffiniPure goat anti-chicken IgY++ (1:400, 103-165-155, Jackson Labs). cFos: Primary, rabbit anti-cFos (1:100, 2250S, cell signaling). Secondary, Alexa Fluor 488-conjugated donkey anti-rabbit IgG (H + L) (1:400, 711-546-152, Jackson Labs).

### Imaging

Confocal images were taken with the Leica TCS SP8. Tile scans are assisted with setting up a tiling experiment with Leica Application Suite software.

### Isoflurane Arousal Test

Optogenetic manipulation mice were anesthetized with 3% isoflurane for 1 minute in an induction chamber. After being quickly attached to the instruments, the mice were placed in a clear isoflurane chamber with optical fibers exiting through a sealed port. After 30 minutes of acclimation, the chamber was equilibrated with 3% isoflurane for 1 minute, then changed to 0.75% isoflurane for 3 minutes before a 20 Hz, 20 ms light pulse (473 or 635 nm) was applied to the animals for 30 seconds. The isoflurane was continued for another 1 minute after the stimulation and switched to 100% O2 to let the mice recover. The time from the laser on to the point when the mice started to wake up (moving, kicking, or tail rising) was recorded as the latency to wake. The trials were repeated three times on the same day from 9:00-18:00. The control group mice received the same treatments. All trials were age-matched with similar animals. All anesthesia chambers were placed on a temperature-controlled heating pad to regulate the temperature between 35–37°C automatically.

### Isoflurane Induction Test

Mice were connected with the photometry system and placed in a clear isoflurane chamber with optical fibers exiting through a sealed port. Photometry data were collected 10 minutes before the induction test as a baseline. Mice were treated with 20 Hz, 20 ms light pulse (635 nm), 1s on, 1s off, from 5 minutes before the induction test to the end. Mice were treated with 3% isoflurane for 3 minutes. The time from the onset time of isoflurane to the point when the mice lost of righting reflex was recorded as LoRR. The trials were conducted from 9:00-18:00 and repeated two times with 1-day intervals. All trials were age-matched with similar animals. All anesthesia chambers were placed on a temperature-controlled heating pad to regulate the temperature between 35–37°C automatically.

### Isoflurane Emergence Test

Mice received 2% isoflurane for 30 minutes in the induction chamber. Afterward, the animals were quickly attached with optical fibers and treated with or without optical stimulation (20 Hz, 20 ms pulse width, 1s on, 1s off) in a clear acrylic chamber open to the room air while placed on their backs to test how quickly the righting reflex can return (RoRR). The RoRR, when the animal turned around and stood on all four paws, was used as the indicator for the emergence from anesthesia. The emergence time from turning off the isoflurane to the RoRR was recorded for analysis. The trials were performed from 9:00 to 18:00 and repeated three times at 3-day intervals. The isoflurane induction chamber was placed on a temperature-controlled heating pad. A temperature probe connected to a temperature controller was placed underneath the animal body to automatically regulate the temperature between 35–37°C. The control group mice received the same treatments. All trials were age-matched with similar animals.

### Hot Plate Test

Mice were anesthetized in the induction chamber with 3% isoflurane for 1 minute to facilitate optical fiber attachment. Then mice went through 5 minutes of optical stimulation with or without the laser (20 Hz, 20 ms pulse width, 1s on, 1s off), followed by the hot plate test. An acrylic container was used as an enclosure to prevent the animal from escaping. The temperature used in the hot plate test was 55°C. Mice were placed onto the hot plate with continuing the optical stimulation until the first time the animals showed licking, fanning, or jumping, at which point the mice were immediately removed from the hot plate. The paw withdrawal latency was recorded for analysis. Mice were removed if there was no response within 45 seconds. The trials were repeated three times on the same day from 9:00 to 15:00, with 60-minute intervals, and again repeated two more times with 3-day intervals. The control group mice received the same treatments. All trials were age-matched with similar animals.

### Formalin Test

With 3% isoflurane for 1 minute of anesthesia, mice were injected with 10 μL of 5% formalin or saline under the skin of the dorsal surface of the left hind paw. Then we placed the mice onto the recording platform, where each animal was individually placed in an acrylic cylinder with a mirror underneath. Then mice were treated with 60 min of optical stimulation with the laser (20 Hz, 20 ms pulse width, 1s on, 1s off). The behaviors were videotaped for 60 min since the mice were placed in the recording platform. The video was analyzed later, and the licking time of the left hind paw was recorded. 0-15 min was considered as the acute phase and 15-60 min was considered as the chronic phase.

### Electrophysiology

The 8-12 weeks Orexin-Cre mice were anesthetized with isoflurane and transcardially perfused with the ice-cold cutting solution containing: 89.5 mM Choline chloride, 20.6 mM Tris, 2.5 mM KCl, 1.2 mM NaH2PO4·2H2O, 20 mM HEPES, 5 mM sodium ascorbate, 2 mM thiourea, 3 mM sodium pyruvate, 25 mM glucose, 30 mM NaHCO3, 10 mM MgSO4·7H2O, 0.5 mM CaCl2·2H2O, 300–310 mOsm, and adjusted to pH 7.4 with HCl. The solution was bubbled with carbogen (95% O2/5% CO2). After we decapitated the mouse and collected the brain. The brain was then cut into 250 μm thick sections with a vibratome (Leica VT1000S). The brain slices were recovered in cutting solution for 10-15 min at 34 °C, then incubated in the HEPES holding artificial cerebrospinal fluid (aCSF) recovery solution, which consisted of 92 mM NaCl, 2.5 mM KCl, 1.25 mM NaH2PO4, 30 mM NaHCO3, 20 mM HEPES, 25 mM glucose, 2 mM thiourea, 5 mM Na-ascorbate, 3 mM Na-pyruvate, 2 mM CaCl2·4H2O and 2 mM MgSO4·7H2O, 300–310 mOsm, and adjusted to pH 7.3-7.4 with HCl. After incubation for 45-60 min at room temperature, we transferred the brain slice to the recording chamber and perfused (3 ml min−1) with the recording aCSF, which consisted of 124 mM NaCl, 2.5 mM KCl, 1.2 mM NaH2PO4·2H2O, 24 mM NaHCO3, 5 mM HEPES, 12.5 mM glucose, 2 mM CaCl2·2H2O and 2 mM MgSO4·7H2O, 300–310 mOsm, and adjusted with NaOH to pH 7.3-7.4. All external solutions were saturated with 95% O2, 5% CO2.

The recording glass pipettes (BF150-86-10, Sutter Instrument) were pulled by a micropipette puller (P1000, Sutter Instrument) into a recording electrode (3–5 MΩ). The recording electrode was filled with a potassium-based internal solution containing 145 mM potassium gluconate, 10 mM HEPES, 1 mM EGTA, 2 mM Mg-ATP, 0.3 mM Na2-GTP, and 2 mM MgCl2 adjusted with KOH to pH 7.2-7.3, 290-300 mOsm. Neurons were identified and subjected to electrophysiological recordings with a 60x water-immersion lens (Zeiss). The recordings were performed using whole-cell techniques (MultiClamp 700B Amplifier, Digidata 1322A analog-to-digital converter) and pClamp 9.2 software (Axon Instruments/Molecular Devices). The traces were digitized at 10kHz. The data were analyzed using pClamp/Clampfit.

Optogenetic stimulation was performed using a 635 nm laser (Intelligent Optogenetics System IOS-635, RWD Life Science Inc). 20 Hz and 20 ms light pulse were given to every trial, five trials per neuron. After breaking into the neurons, the neurons were held at −65 mV to record EPSCs, and 0 mV to record IPSCs. Voltage-clamp experiments were carried out to record EPSCs and IPSCs, and current clamp tests were performed to record action potentials. 10 μM SB-334867(SML1530, Millipore Sigma), and 30 μM TCS OX2 29 HCl (SML2879, Millipore Sigma) were added into aCSF perfusions to block the orexin 1 receptor, and orexin 2 receptor separately.

### RNAscope in situ hybridization

C57/BL6 mice (14-16 weeks) were transcardially perfused as described above in the Immunohistochemistry section. Brains were post-fixed at 4°C in 30% sucrose for 24 hours and cut into 15 μm. Slides were stored at −80°C until staining. The staining was performed with the RNAscope Multiplex Fluorescent Detection Reagents v2 kit (323110, Advanced Cell Diagnostics) according to the user manual. Probes for OX1R (Hcrtr1, Cat No. 466631-C1), ChAT (CHAT, Cat No. 408731-C1), vGlut2 (Slc17a6, Cat No. 428871-C2), vGAT (Slc32a1, Cat No. 319191-C2), TH (TH, Cat No. 317621-C2), and OX2R (Hcrtr2, Cat No. 581631-C3) were commercially available by Advanced Cell Diagnostics.

In brief, slides were post-fixed in 4% PFA for 2 hours and briefly washed in PBS. After being treated with hydrogen peroxide (H2O2) for 10 min at room temperature, washed in diethylpyrocarbonate (DEPC)-treated Millipore water (ddH2O), and boiled in Target Retrieval solution (98-102 °C) for 5 min. After a brief washing in water and dehydration in absolute ethanol, dried at 60 °C incubator for 5 min. Moisten the slide with ddH2O, slides were incubated with protease Ill for 15 min at 40°C in the HybEZ oven (Advanced Cell Diagnostics). Slides were washed again in ddH2O and hybridized with the mixture of probes in different channels for 2 hours in a HybEZ oven at 40°C. Afterward, the hybridization was amplified with AMP-1 for 30min, AMP-2 for 30 min, and AMP-3 for 15 min, respectively. The signals were then developed for each channel by incubating with HRP-C1 for 15 min, the Opal™ 520 Reagent Pack (FP1487001KT, Akoya Biosciences Inc) for 30min, and HRP blocker for 15 min. HRP-C2 for 15 min, the Opal™ 620 Reagent Pack (FP1495001KT, Akoya Biosciences Inc) for 30 min, and HRP blocker for 15 min. HRP-C3 for 15 min, the Opal™ 690 Reagent Pack (FP1497001KT, Akoya Biosciences Inc) for 30 min, and the HRP blocker for 15 min. All amplification and development were performed at 40 °C in the HybEZ oven, and the slides were washed using an ACD wash buffer after each step. After that, the slides were mounted with DAPI Fluoromount-G mounting medium (0100-20, SouthernBiotech), and covered with coverslips. Imaging was performed with a confocal laser scanning microscope Leica TCS SP8 with a 40x oil immersion objective. The RNAscope images were manually adjusted with ImageJ/Fiji and the data were automatically analyzed with the QuPath V0.5.1.

### Data analysis and statistics

Electrophysiology data was analyzed with Clampfit 10 (Molecular Devices). EPSC and IPSC amplitudes were calculated as the difference between the peak amplitude in a pre-defined window after the light stimulation onset and the mean amplitude just preceding the EPSC or IPSC. EPSC or IPSC latency was measured as the time from the onset of laser stimulation to the first intersection between the baseline and the EPSC or IPSC, which can be easily identified at the place of maximal rising/falling curvature. EEG data was analyzed with MATLAB (R2023B; MathWorks). Fiber photometry data was processed with the RWD fiber photometry software. The figures were created using Prism 10.3.1(GraphPad Software Inc. San Diego, CA, USA), MATLAB, ImageJ, QuPath, and Illustrator (Adobe).

Statistical details are presented in the figure legends. All data were analyzed using Prism 10.3.1. Two-tailed, unpaired t-test was used to analyze the significance between two groups with normally distributed data. Ordinary One-way ANOVA was used to compare the significance of three or more groups with one independent variable. Two-way ANOVA was used to compare the mean differences between groups with two independent variables. ANOVAs were followed by post-hoc tests with multiple comparison corrections, as indicated in the figure legends. All data are shown as mean ± standard error of the mean (S.E.M.).

## Details of authors’ contributions

W.Z. and X.X. designed and performed research; F.W. contributed to the electrophysiology experimental design and data analysis; C.C contributed to virus production and RNAscope design; Z.G. contributed as scientific advisor; W.Z. contributed to the project administration and funding acquisition; W.Z. and X.X. wrote the paper.

## Acknowledgments

We thank Drs. Lily Jan and Yuh-Nung Jan for sharing the lab resources; Dr. Akihiro Yamanaka for sharing the frozen embryo of the orexin-Cre mouse line; Tong Cheng, Marena Tynan-La Fontaine for their technical support; Dr. Yanmeng Guo, Dr. Wendy W.S. Yue, and Yuzhang Chen for insightful discussions. All experimental data were generated at the University of California, San Francisco.

## Data availability

All data generated in this study are included within the main text or supplementary materials, with source data provided. This includes individual data points and averaged values presented in both the figures and supplementary information. The corresponding authors can provide raw data from optogenetic stimulation, in vivo fiber photometry, in vitro electrophysiology, and other experiments upon request.

## Supplementary data

**Supplementary Fig. 1.**
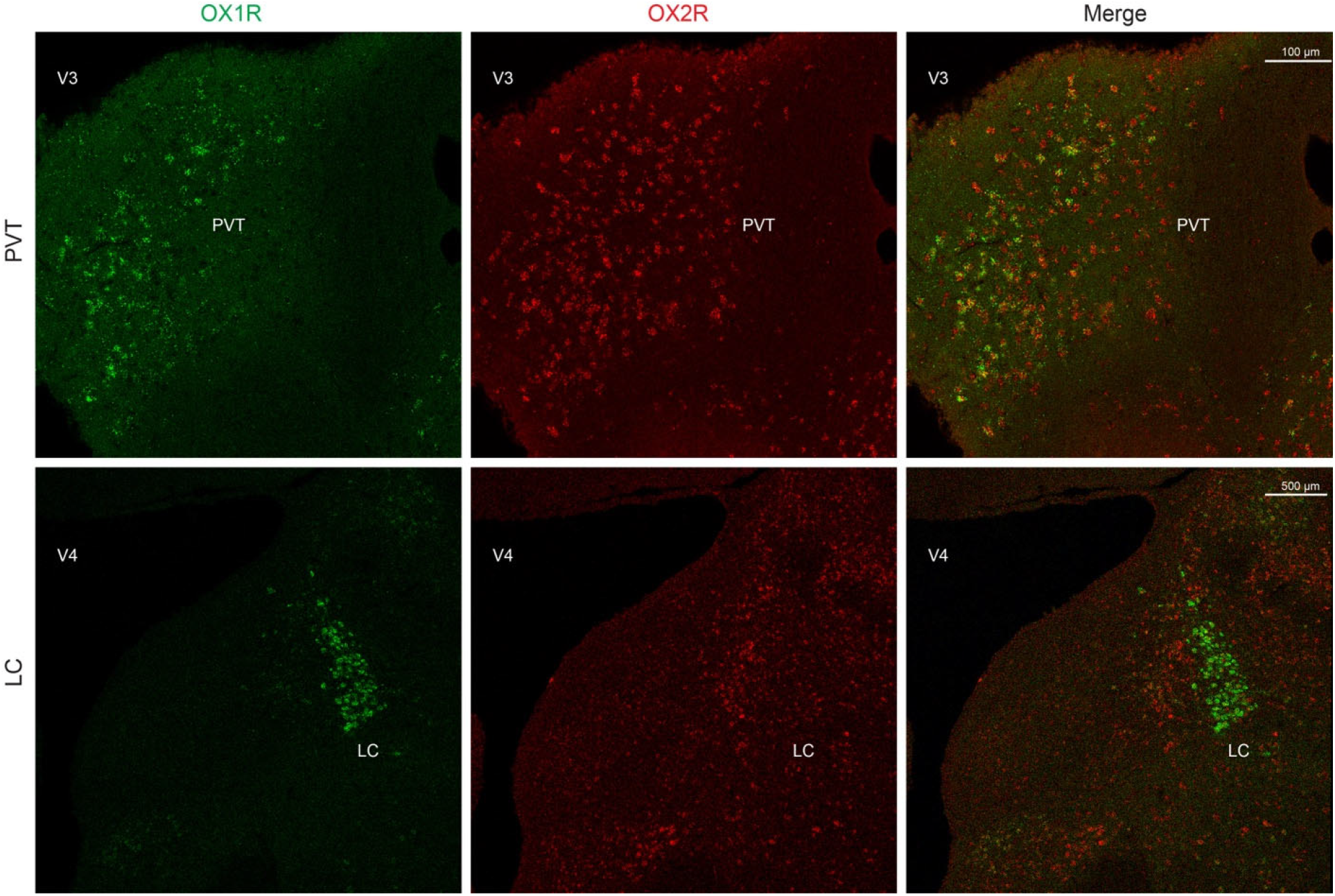
OX1R and OX2R distributions at different regions. RNAscope staining of OX1R and OX2R at the paraventricular nucleus of the Thalamus (PVT) and the locus coeruleus (LC) shows the predominant OX1R staining in the LC but evenly distributed OX1R and OX2R in the PVT.

**Supplementary Fig. 2.**
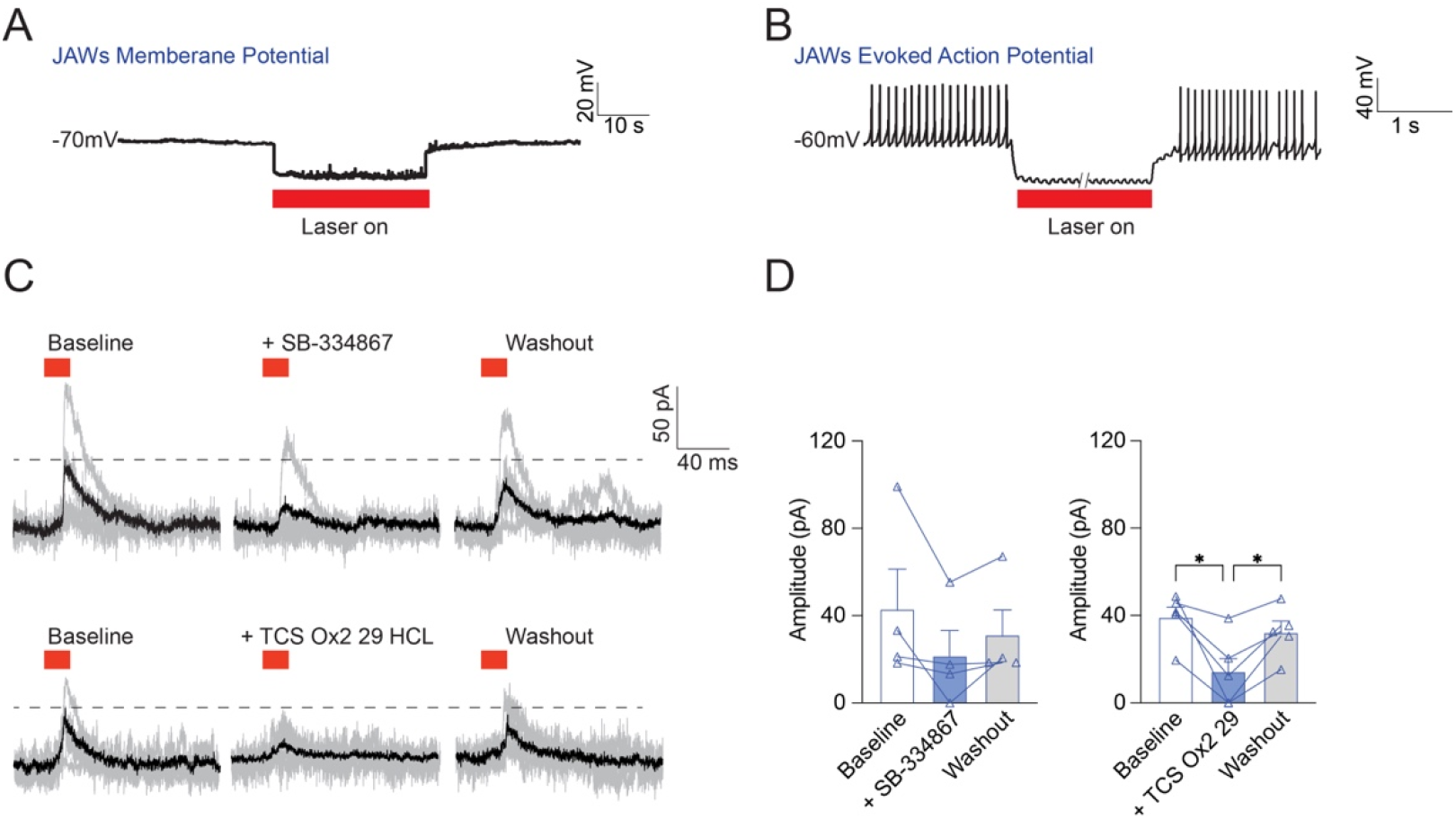
Acute brain slice electrophysiology recordings. **A, B**. *AAV5-FLEX-JAWs-GFP* was injected into the LHA of *Orexin-Cre* mice. Patch-clamp whole-cell recordings on LHA orexin neurons showed optogenetic activation of JAWs decreased the membrane potential (A) and silenced the action potentials evoked by injecting 50 pA current at 10 Hz (B). **C**. The IPSCs can be blocked by the OX2R antagonist - TCS OX2 29, but not as much by the OX1R antagonist - SB-334867. **D**. The amplitudes before, during, and after blockage. One-way ANOVA, the effect of the SB-334867SB-334867, F_(1.174, 3.521)_ = 3.93, p = 0.128, Tukey’s multiple comparisons of the baseline (43 ± 19.01 pA), SB-334867 (21.6 ± 11.86 pA), and washout (31.26 ± 11.97 pA) showed no significant difference (baseline vs SB-334867: p = 0.2351, SB-334867 vs washout: p = 0.2105); n = 4. One-way ANOVA, the effect of the TCS OX2, F_(1.067, 4.269)_ = 13.45, p = 0.0186, Tukey’s multiple comparisons of the baseline (39.2 ± 5.13 pA), TCS OX2 (14.32 ± 7.23 pA), and washout (32.28 ± 5.19 pA) showed TCS OX2 29 significantly decreased the amplitude of the laser evoked IPSCs (baseline vs TCS OX2 29: p = 0.0478, TCS OX2 vs washout: p = 0.0218); n = 5. n stands for cell numbers, each data point represents the average of five traces from a single cell. *p ≤ 0.05. *p ≤ 0.05. All data are expressed as mean ± S.E.M.

**Supplementary Fig. 3.**
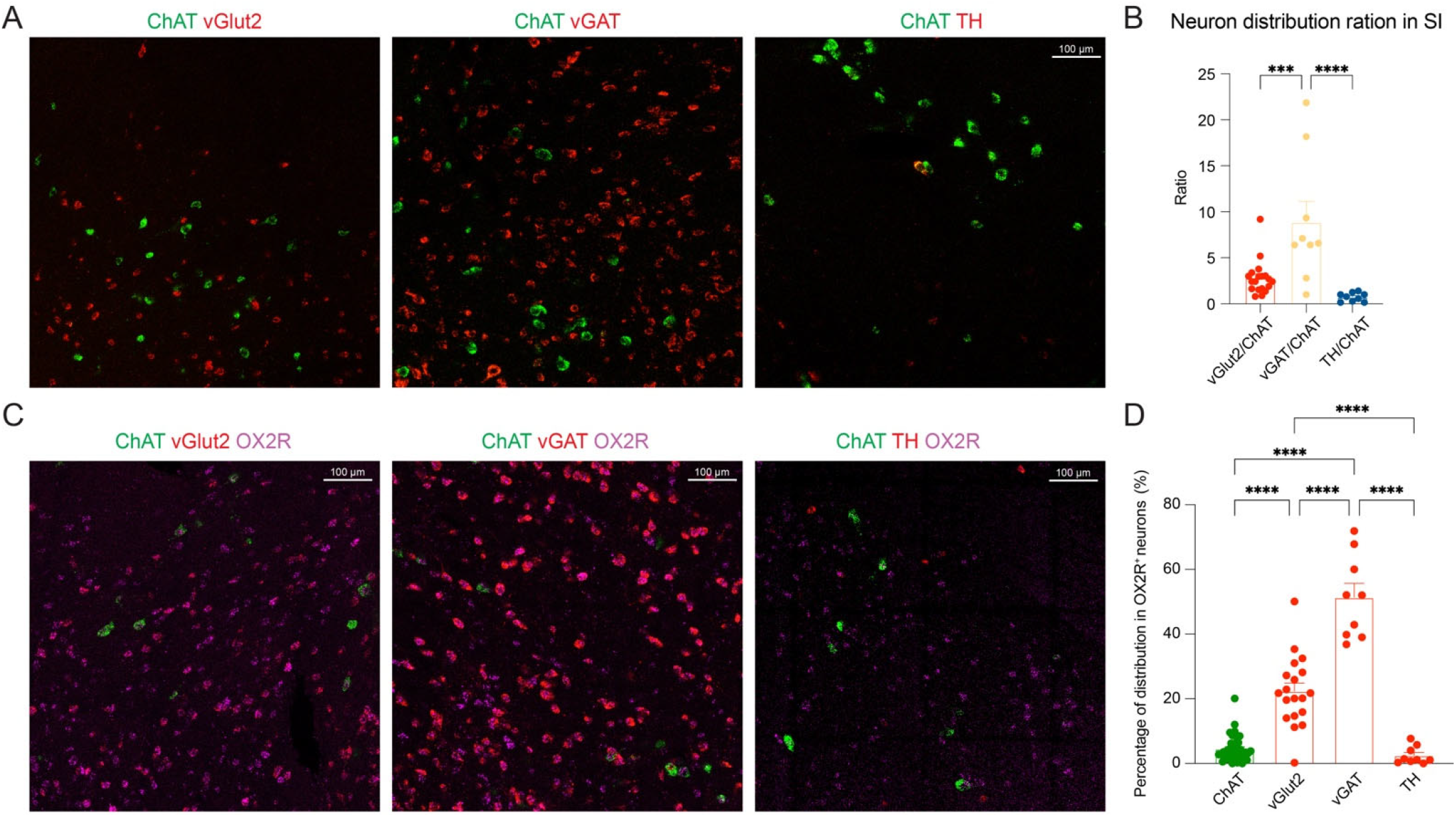
Colocalization of OX2R, vGlut2, vGAT, ChAT, and TH in the SI. **A**. Double-staining of vGlut2/ChAT, vGAT/ChAT, and TH/ChAT. **B**. Ratios of vGlut2/ChAT, vGAT/ChAT, and TH/ChAT. Ordinary one-way ANOVA, F_(2, 34)_ = 12.9, p < 0.0001), Šidák’s multiple comparisons showed the ratio of vGAT/ChAT (8.86 ± 2.28, n = 9 brain slices from 5 mice) was significantly higher than vGlut2/ChAT (2.81 ± 0.43, p = 0.0006, n = 19 brain slices from 5 mice) and TH/ChAT (0.73 ± 0.16, p < 0.0001, n = 9 brain slices from 5 mice). **C**. The co-localization of ChAT, vGlut2, vGAT, TH, and OX2R. **D**. The percentage of ChAT, vGlut2, vGAT, and TH neurons in the OX2R+ neurons. Data was calculated from the number of double positive neurons divided by the total number of OX2R+ neurons. Ordinary one-way ANOVA, F_(3, 70)_ = 105.2, p < 0.0001. Tukey’s multiple comparisons showed vGlut2 (22.34 ± 2.48 %, n = 19 brain slices from 5 mice), vGAT (51.38 ± 4.31 %, n = 9 brain slices from 5 mice) ChAT (4.53 ± 0.66 %, n = 37 brain slices from 5 mice), and TH (2.47 ± 0.9 %, n = 9 brain slices from 5 mice) are different from each other. ****p < 0.0001. All data are expressed as mean ± S.E.M.

## Reference

1. Steen, P. A. & Michenfelder, J. D. Neurotoxicity of anesthetics. Anesthesiology 50, 437–453 (1979).

2. Neukirchen, M. & Kienbaum, P. Sympathetic nervous system: evaluation and importance for clinical general anesthesia. Anesthesiology 109, 1113–31 (2008).

3. Mashour, G. A. et al. Recovery of consciousness and cognition after general anesthesia in humans. eLife 10, e59525 (2021).

4. Moody, O. A. et al. The Neural Circuits Underlying General Anesthesia and Sleep. Anesth. Analg. 132, 1254–1264 (2021).

5. Sakurai, T. et al. Orexins and orexin receptors: a family of hypothalamic neuropeptides and G protein-coupled receptors that regulate feeding behavior. Cell 92, 573–85 (1998).

6. de Lecea, L. et al. The hypocretins: hypothalamus-specific peptides with neuroexcitatory activity. Proc. Natl. Acad. Sci. U. S. A. 95, 322– 327 (1998).

7. Tyree, S. M. et al. Optogenetic and pharmacological interventions link hypocretin neurons to impulsivity in mice. Commun. Biol. 6, 1–8 (2023).

8. Inutsuka, A. et al. Concurrent and robust regulation of feeding behaviors and metabolism by orexin neurons. Neuropharmacology 85, 451–60 (2014).

9. Elam, H. B., Perez, S. M., Donegan, J. J. & Lodge, D. J. Orexin receptor antagonists reverse aberrant dopamine neuron activity and related behaviors in a rodent model of stress-induced psychosis. Transl. Psychiatry 11, 1–11 (2021).

10. Soya, S. et al. Orexin Receptor-1 in the Locus Coeruleus Plays an Important Role in Cue-Dependent Fear Memory Consolidation. J. Neurosci. 33, 14549–14557 (2013).

11. Lin, L. et al. The sleep disorder canine narcolepsy is caused by a mutation in the hypocretin (orexin) receptor 2 gene. Cell 98, 365–76 (1999).

12. Ito, H. et al. Deficiency of orexin signaling during sleep is involved in abnormal REM sleep architecture in narcolepsy. Proc. Natl. Acad. Sci. 120, e2301951120 (2023).

13. Chemelli, R. M. et al. Narcolepsy in orexin knockout mice: molecular genetics of sleep regulation. Cell 98, 437–51 (1999).

14. Flanigan, M. E. et al. Orexin signaling in GABAergic lateral habenula neurons modulates aggressive behavior in male mice. Nat. Neurosci. 23, 638–650 (2020).

15. Tung, L.-W. et al. Orexins contribute to restraint stress-induced cocaine relapse by endocannabinoid-mediated disinhibition of dopaminergic neurons. Nat. Commun. 7, 12199 (2016).

16. Date, Y. et al. Orexins, orexigenic hypothalamic peptides, interact with autonomic, neuroendocrine and neuroregulatory systems. Proc Natl Acad Sci U A 96, 748–53 (1999).

17. Bulbul, M., Babygirija, R., Ludwig, K. & Takahashi, T. Central orexin-A increases gastric motility in rats. Peptides 31, 2118–22 (2010).

18. Varga, A. G., Whitaker-Fornek, J. R., Maletz, S. N. & Levitt, E. S. Activation of orexin-2 receptors in the Kölliker-Fuse nucleus of anesthetized mice leads to transient slowing of respiratory rate. Front. Physiol. 13, 977569 (2022).

19. Li, J. et al. Orexin activated emergence from isoflurane anaesthesia involves excitation of ventral tegmental area dopaminergic neurones in rats. Br. J. Anaesth. 123, 497–505 (2019).

20. Inutsuka, A. et al. The integrative role of orexin/hypocretin neurons in nociceptive perception and analgesic regulation. Sci Rep 6, 29480 (2016).

21. Suzuki, M., Shiraishi, E., Cronican, J. & Kimura, H. Effects of the orexin receptor 2 agonist danavorexton on emergence from general anaesthesia and opioid-induced sedation, respiratory depression, and analgesia in rats and monkeys. Br. J. Anaesth. 132, 541–552 (2024).

22. Yan, J.-A. et al. Orexin affects dorsal root ganglion neurons: a mechanism for regulating the spinal nociceptive processing. Physiol. Res. 57, 797–800 (2008).

23. Azhdari-Zarmehri, H., Semnanian, S. & Fathollahi, Y. Orexin-A microinjection into the rostral ventromedial medulla causes antinociception on formalin test. Pharmacol. Biochem. Behav. 122, 286–290 (2014).

24. Shakerinava, P., Sayarnezhad, A., Karimi-Haghighi, S., Mesgar, S. & Haghparast, A. Antagonism of the orexin receptors in the ventral tegmental area diminished the stress-induced analgesia in persistent inflammatory pain. Neuropeptides 96, 102291 (2022).

25. Kaneko, T. et al. Orexin neurons play contrasting roles in itch and pain neural processing via projecting to the periaqueductal gray. Commun. Biol. 7, 290 (2024).

26. Xiang, X. et al. Neuroanatomical Basis for the Orexinergic Modulation of Anesthesia Arousal and Pain Control. Front. Cell. Neurosci. 16, 891631 (2022).

27. Scharf, M. T. & Kelz, M. B. Sleep and Anesthesia Interactions: A Pharmacological Appraisal. Curr Anesth. Rep 3, 1–9 (2013).

28. Ozen Irmak, S. & de Lecea, L. Basal Forebrain Cholinergic Modulation of Sleep Transitions. Sleep 37, 1941–1951 (2014).

29. Zhu, Z. et al. A substantia innominata-midbrain circuit controls a general aggressive response. Neuron 109, 1540-1553.e9 (2021).

30. Jones, B. E. Activity, modulation and role of basal forebrain cholinergic neurons innervating the cerebral cortex. Prog. Brain Res. 145, 157– 169 (2004).

31. Leung, L. S., Chu, L., Prado, M. A. M. & Prado, V. F. Forebrain Acetylcholine Modulates Isoflurane and Ketamine Anesthesia in Adult Mice. Anesthesiology 134, 588–606 (2021).

32. Xu, M. et al. Basal Forebrain Circuit for Sleep-Wake Control. Nat. Neurosci. 18, 1641–1647 (2015).

33. Cui, Y. et al. A Central Amygdala-Substantia Innominata Neural Circuitry Encodes Aversive Reinforcement Signals. Cell Rep. 21, 1770–1782 (2017).

34. Dafny, N. et al. Lateral hypothalamus: Site involved in pain modulation. Neuroscience 70, 449–460 (1996).

35. Wang, D. et al. Lateral septum-lateral hypothalamus circuit dysfunction in comorbid pain and anxiety. Mol. Psychiatry 28, 1090– 1100 (2023).

36. Duffet, L. et al. A genetically encoded sensor for in vivo imaging of orexin neuropeptides. Nat. Methods 19, 231–241 (2022).

37. Klapoetke, N. C. et al. Independent Optical Excitation of Distinct Neural Populations. Nat. Methods 11, 338–346 (2014).

38. Chuong, A. S. et al. Noninvasive optical inhibition with a red-shifted microbial rhodopsin. Nat. Neurosci. 17, 1123–1129 (2014).

39. Bingham, S. et al. Orexin-A, an hypothalamic peptide with analgesic properties. Pain 92, 81–90 (2001).

40. Holloway, B. B. et al. Monosynaptic Glutamatergic Activation of Locus Coeruleus and Other Lower Brainstem Noradrenergic Neurons by the C1 Cells in Mice. J. Neurosci. 33, 18792–18805 (2013).

41. Munn, B. R., Müller, E. J., Wainstein, G. & Shine, J. M. The ascending arousal system shapes neural dynamics to mediate awareness of cognitive states. Nat. Commun. 12, 6016 (2021).

42. Jones, B. E. Arousal and sleep circuits. Neuropsychopharmacology 45, 6–20 (2020).

43. Gouras, G. K., Rance, N. E., Scott Young, W. & Koliatsos, V. E. Tyrosine-hydroxylase-containing neurons in the primate basal forebrain magnocellular complex. Brain Res. 584, 287–293 (1992).

44. Trivedi, P., Yu, H., MacNeil, D. J., Van der Ploeg, L. H. & Guan, X. M. Distribution of orexin receptor mRNA in the rat brain. FEBS Lett 438, 71–5 (1998).

45. Soya, S. et al. Orexin modulates behavioral fear expression through the locus coeruleus. Nat Commun 8, 1606 (2017).

46. González, J. A., Jensen, L. T., Fugger, L. & Burdakov, D. Convergent inputs from electrically and topographically distinct orexin cells to locus coeruleus and ventral tegmental area. Eur. J. Neurosci. 35, 1426– 1432 (2012).

47. Manns, I. D., Alonso, A. & Jones, B. E. Discharge Profiles of Juxtacellularly Labeled and Immunohistochemically Identified GABAergic Basal Forebrain Neurons Recorded in Association with the Electroencephalogram in Anesthetized Rats. J. Neurosci. 20, 9252– 9263 (2000).

48. Gritti, I., Mainville, L. & Barbara, J. Codistribution of GABA-with acetylcholine-synthesizing neurons in the basal forebrain of the rat. https://onlinelibrary.wiley.com/doi/abs/10.1002/cne.903290403?sid=nlm%3Apubmed (1993).

49. Yang, C., Thankachan, S., McCarley, R. W. & Brown, R. E. The menagerie of the basal forebrain: How many (neural) species are there, what do they look like, how do they behave and who talks to whom? Curr. Opin. Neurobiol. 44, 159–166 (2017).

50. Cardin, J. A. et al. Driving fast-spiking cells induces gamma rhythm and controls sensory responses. Nature 459, 663–667 (2009).

51. Buzsaki, G. et al. Nucleus basalis and thalamic control of neocortical activity in the freely moving rat. J. Neurosci. 8, 4007–4026 (1988).

52. Záborszky, L. et al. Specific Basal Forebrain–Cortical Cholinergic Circuits Coordinate Cognitive Operations. J. Neurosci. 38, 9446–9458 (2018).

53. Peng, W. et al. Regulation of sleep homeostasis mediator adenosine by basal forebrain glutamatergic neurons. Science 369, eabb0556 (2020).

54. Agostinelli, L. J. et al. Descending Projections from the Basal Forebrain to the Orexin Neurons in Mice. J. Comp. Neurol. 525, 1668– 1684 (2017).

55. Razavi, B. M. & Hosseinzadeh, H. A review of the role of orexin system in pain modulation. Biomed. Pharmacother. 90, 187–193 (2017).

56. Lee, H.-J. et al. Stress induces analgesia via orexin 1 receptor-initiated endocannabinoid/CB1 signaling in the mouse periaqueductal gray. Neuropharmacology 105, 577–586 (2016).

57. Ho, Y.-C. et al. Activation of Orexin 1 Receptors in the Periaqueductal Gray of Male Rats Leads to Antinociception via Retrograde Endocannabinoid (2-Arachidonoylglycerol)-Induced Disinhibition. J. Neurosci. 31, 14600–14610 (2011).

58. Emam, A. H. et al. Modulation of nociception by medial pre-optic area orexin a receptors and its relation with morphine in male rats. Brain Res Bull 127, 141–147 (2016).

59. Yazdi-Ravandi, S., Razavi, Y., Haghparast, A., Goudarzvand, M. & Haghparast, A. Orexin A induced antinociception in the ventral tegmental area involves D1 and D2 receptors in the nucleus accumbens. Pharmacol. Biochem. Behav. 126, 1–6 (2014).

